# Chemical-induced Gene Expression Ranking and its Application to Pancreatic Cancer Drug Repurposing

**DOI:** 10.1101/2021.12.13.472490

**Authors:** Thai-Hoang Pham, Yue Qiu, Jiahui Liu, Steven Zimmer, Eric O’Neill, Lei Xie, Ping Zhang

## Abstract

Chemical-induced gene expression profiles provide critical information on the mode of action, off-target effect, and cellar heterogeneity of chemical actions in a biological system, thus offer new opportunities for drug discovery, system pharmacology, and precision medicine. Despite their successful applications in drug repurposing, large-scale analysis that leverages these profiles is limited by sparseness and low throughput of the data. Several methods have been proposed to predict missing values in gene expression data. However, most of them focused on imputation and classification settings which have limited applications to real-world scenarios of drug discovery. Therefore, a new deep learning framework named chemical-induced gene expression ranking (CIGER) is proposed to target a more realistic but more challenging setting in which the model predicts the rankings of genes in the whole gene expression profiles induced by *de novo* chemicals. The experimental results show that CIGER significantly outperforms existing methods in both ranking and classification metrics for this prediction task. Furthermore, a new drug screening pipeline based on CIGER is proposed to select approved or investigational drugs for the potential treatments of pancreatic cancer. Our predictions have been validated by experiments, thereby showing the effectiveness of CIGER for phenotypic compound screening of precision drug discovery in practice.

## Introduction

Phenotypic screening has been shown to be more effective than target-based screening for first-in-class drug discovery but this approach also has some limitations due to the low-throughput of phenotypic assays^1^. Recently, several high-throughput phenotypic datasets that cover the wide ranges of chemical compounds and cell lines have been developed to alleviate this problem. Gene expression profiling method based on these datasets has been shown to be a very effective and powerful tool for phenotypic drug discovery and system pharmacology. Computational techniques that leverage genome-wide gene expression, especially chemical-induced differential gene expression, has demonstrated a great potential in drug repurposing^2–5^, elucidation of drug mechanisms^6^, lead identification^7^, and predicting side effect of drug compounds^8^.

Pioneered by Connectivity Map^9^, a database that consists of ~ 1, 300 chemical-induced gene expression profiles of five human cancer cell lines, many works have been proposed to identify existing drugs for the treatment of new diseases by selecting drugs that reverse the disease gene expressions^10,11^. However, the low coverage across cell types in Connectivity Map limited the performances of those methods, especially in large-scale analysis settings. To alleviate this limitation, a novel and affordable gene expression profiling method has been proposed. In particular, Library of Integrated Network-based Cell-Signature (LINCS) program introduced L1000 platform that measured the expression of the most informative genes (i.e., ~1, 000 landmark genes) instead of whole-genome data, thus reducing the cost for measuring each gene expression profile to ~ $5^12^. This profiling technique resulted in a gene expression dataset, called LINCS L1000, which consists of ~ 1, 400, 000 gene expression profiles covering the responses of ~ 20, 000 compounds at different concentrations across ~ 80 human cell lines. Despite the significantly increasing coverage of compounds and cell lines in L1000, large-scale analysis based on this dataset is still limited due to several problems. First, in spite of the wide coverage across cell lines, compounds, and concentrations, there are many missing expression values in the vast and high-dimensional combinatorial space of chemicals, concentrations, and cell lines. Moreover, there are hundreds of millions of drug-like chemicals so it is not feasible to measure gene expression profiles across a large number of cell lines for all of these chemicals. Second, LINCS L1000 and other gene expression datasets are highly noisy due to experimental limitations^13,14^. As a result, many experiment measurements are not reliable in these datasets. These problems seriously affect the performances of large-scale genome analysis using LINCS L1000, and motivate the development of computational methods to predict missing gene expression values in this high-dimensional combinatorial space.

Several works have been proposed to predict gene expression values for chemical-induced gene expression data in general^15–22^, and for LINCS L1000 in particular^13,23,24^ but most of them focused on the imputation and classification settings only. In particular, they predict either expression values or classes of certain genes in the gene expression profiles or the whole gene expression profiles of certain existing chemicals. The imputation setting is not practical and useful in the real-world application of drug discovery, in which the assessment about novel chemicals (i.e., chemicals not in gene expression dataset) can not be made due to the unavailability of the corresponding gene expression profiles. Moreover, formulating this problem as a classification problem has limited scope for practical applications because this setting focuses only on a small subset of genes while down-stream analysis based on gene expression profiles often benefits most when using the information of the whole profiles (i.e., ranking of genes)^25^. redThere have been also some works proposed in recommender system context for predicting the ranking of items in data^26,27^. However, these methods are designed for matrix data only, and hence they cannot be adapted to work with LINCS L1000 dataset which is formulated as high-dimensional data.

In this work, we propose a new framework named chemical-induced gene expression ranking (CIGER) that can predict gene ranking in L1000 gene expression profiles induced by *de novo* chemicals. In particular, CIGER is a neural network-based architecture that leverages the representations of biological objects including chemicals, cell lines, and genes to predict the gene ranking in the corresponding gene expression profiles. This framework consists of several components as follows. First, due to the importance of ranking information with respect to gene expressions^25^, we focus on the prediction on the whole gene expression profile by using some ranking loss functions^28–33^ instead of considering prediction on each gene separately by some regression or classification loss functions in the optimization process. Second, we learn the contextualized representations for genes before making predictions by using an attention mechanism named multi-head attention^34^ to capture the dependencies among genes, chemicals, and cell lines. We also utilize graph convolutional network^35^ to extract useful information from the graph structure of chemicals. Finally, the multi-layer feed-forward neural network is used to predict gene ranking from the contextualized representations. Figure 1 presents the overall architecture of CIGER and the details of this model are shown in Experimental Procedures section. We evaluate the effectiveness of CIGER for predicting gene expression ranking and classification tasks on LINCS L1000 dataset under a 5-fold cross-validation setting. The results show that CIGER significantly outperforms other models across all ranking and classification metrics. Furthermore, we design a new *in silico* drug screening pipeline for finding potential treatments from all drugs in the DrugBank database for pancreatic cancer based on their chemical-induced gene expression profiles (i.e., gene rankings) generated by CIGER. This pipeline demonstrates that CIGER can facilitate phenotypic compound screening for precision drug discovery in practice. In summary, the contributions of this work are as follow:

**Figure 1.**
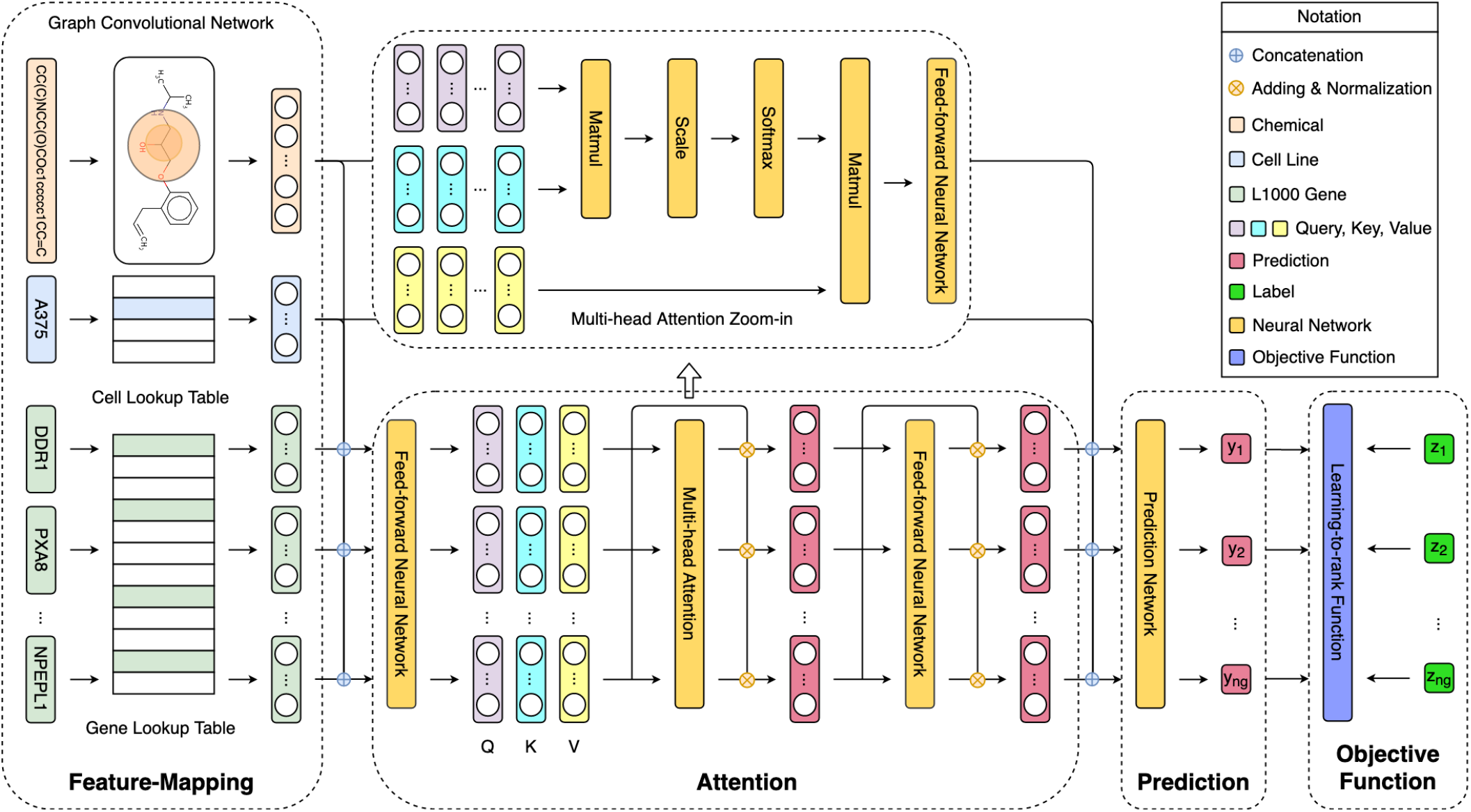
Overview architecture of Chemical-induced Gene Expression Ranking (CIGER). This model consists of the four main components: feature-mapping, attention, prediction, and learning-to-rank objective function. It takes input as a tuple of chemical structure, cell line, L1000 genes and then predicts the ranking of genes in the corresponding gene expression profile. Note that the multi-head attention zoom-in is detailed architecture of the multi-head attention layer in CIGER and is separated from the main figure.

- We propose a deep learning framework (CIGER) that leverages chemical, cell, and gene representations to predict gene ranking in chemical-induced gene expression profiles for *de novo* chemicals which is a more practical but more challenging problem.
- Leveraging CIGER, we design a new phenotypic (i.e., gene expression) drug repurposing pipeline and use pancreatic cancer as a showcase although it can be easily applied for finding treatments for other diseases.
- Source code and the generated gene signatures of all drugs in DrugBank are made available for research purposes at https://github.com/pth1993/CIGER.

## Results and Discussion

### Chemical-induced Gene Expression Data Analysis

Several genome-wide chemical-induced gene expression datasets have been published and applied in drug discovery and system pharmacology, and the LINCS L1000 dataset^12^ is the largest and latest dataset among them. This dataset includes the gene expression profiles generated from a platform called L1000. In specific, this platform measures the expression of 978 landmark genes which captures most of the information from the entire transcriptome. Since the first release of the LINCS L1000 dataset that includes more than 1.3 million gene expression profiles from ~ 20,000 small molecule compounds over 77 cell lines, there have been many works focusing on improving the quality of this dataset^36–38^. In our study, we experiment with an L1000 dataset using Bayesian analysis for calculating peak deconvolution^39^. This dataset has been shown to generate more robust z-score profiles from L1000 assay data compared to the original L1000 dataset using k-means clustering for calculating peak deconvolution^12^, and therefore, gives better representation for chemicals. Initially, we investigate the sparse and noisy problems of this gene expression dataset by calculating Pearson correlation scores among bio-replicate gene expression profiles (level 4 data) of experiments and then visualizing these scores in the chemical-cell line space. From Figure 2, we can observe that only 5.36% of experiments are available in this combinatorial space (i.e., 21229 chemicals × 83 cell lines) and among existing experiments, only 8.47% of them have the corresponding Pearson correlation scores *>* 0.6. These obstacles certainly hinder the utility of this dataset to its downstream applications in drug discovery. Figure 2 also shows the statistics of this dataset with respect to cell lines, exposure times, and chemical concentrations. We can see that the top 10 most popular cell lines account for 77.14% of the number of experiments and the most popular time exposure and chemical concentration are 24 hours (67.56%) and 10 µM (44.82%) respectively.

**Figure 2.**
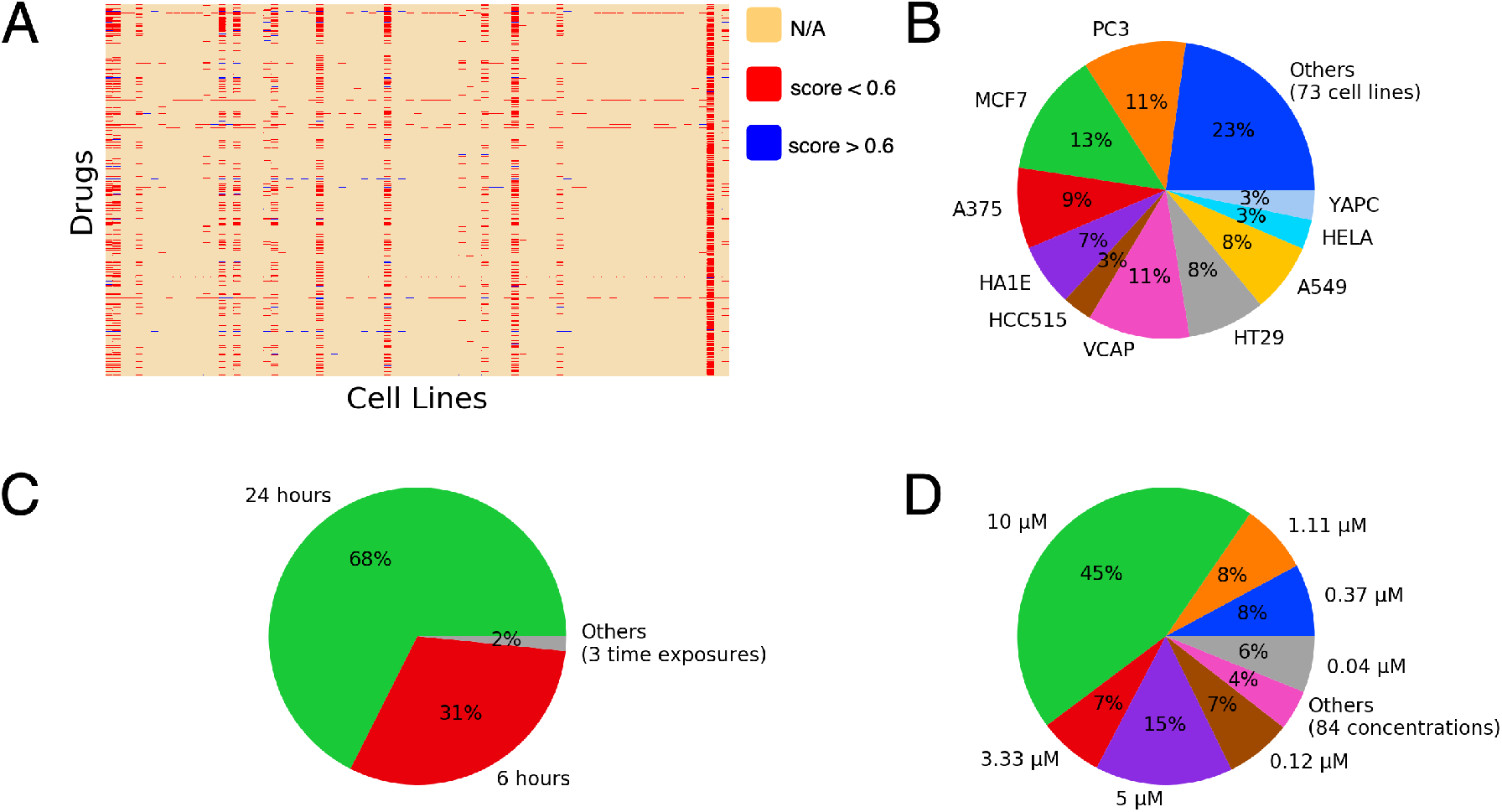
LINCS L1000 data statistical analysis (cell lines, dosages, and time exposures are shown in random order). (A) Gene expression profiles in chemical-cell line space (i.e., yellow color denotes missing profiles for chemical-cell line pairs, red and blue colors denote the pairs with the corresponding correlation scores is smaller (unstable) and larger (stable) than 0.6 respectively). (B) Proportion of profiles by cell lines. (C) Proportion of profiles by time exposures. (D) Proportion of profiles by chemical concentrations.

In our study, to reduce the noise of this dataset, we only select the gene expression profiles (level 5 data) of the 10 most popular cell lines (i.e. A375, A549, HA1E, HCC515, HELA, HT29, MCF7, PC3, VCAP, YAPC) in both phase I (GSE92742) and phase II (GSE70138) of this dataset that satisfy two conditions: (1) the average correlation scores among their bio-replicates (level 4 data) larger than 0.6 and (2) the concentration and exposure time of chemicals are largest (i.e., 10 µM and 24h respectively). The resulting dataset includes some duplicate experiments (i.e., experiments with the same chemical and cell line) so we calculate the ranking of each L1000 gene across duplicate experiments and then select the experiment that has the most genes closed to the median.

Ranking loss functions focus on optimizing the top-ranked objects only while in gene expression analysis, both most up-regulated (positive z-score) and down-regulated (negative z-score) genes are important so we multiply the z-scores in the gene expression profiles with −1 when training the model to rank down-regulated genes. After processing, the data consists of 3,294 gene expression profiles. The number of chemicals and the statistics of gene expression values corresponding to each cell line are shown in Table 1.

**Table 1.** Number of chemicals and gene expression statistics across cell lines for gene expression dataset after processing.

|  | #Gene Expression Profile (3,294) | Gene Expression Value |  |  |
| --- | --- | --- | --- | --- |
|  |  | Max | Mean | Min |
| A375 | 430 | 7.8573 | -0.0161 | -7.6315 |
| A549 | 232 | 6.5863 | 0.0059 | -6.4124 |
| HA1E | 394 | 6.6578 | -0.0100 | -6.6511 |
| HCC515 | 262 | 6.6362 | -0.0027 | -6.1765 |
| HELA | 191 | 5.2012 | -0.0138 | -5.1913 |
| HT29 | 334 | 5.8744 | 0.0179 | -5.7641 |
| MCF7 | 561 | 8.8711 | 0.0067 | -8.7236 |
| PC3 | 481 | 8.6328 | -0.0181 | -8.5835 |
| VCAP | 256 | 6.6322 | -0.0158 | -6.3357 |
| YAPC | 153 | 5.1836 | -0.0504 | -5.1830 |

### Gene Expression Ranking for *de novo* Chemicals

To validate the effectiveness of CIGER for predicting gene expression ranking for novel chemicals, we conduct experiments on the LINCS L1000 dataset^39^ to compare its prediction performances with existing methods including DeepCOP^24^ and Tensor-train Weight Optimization (TT-WOPT)^40^. The detailed architectures of these models are presented in Experimental Section. Because our study focuses on predicting gene expression ranking for novel chemicals, we perform experiments under 5-fold cross-validation (i.e., train : dev : test = 60 : 20 : 20) divided by chemicals to ensure the chemicals in the development and testing sets are not seen in the training set. Normalized Discounted Cumulative Gain (NDCG) and Precision@K (Experimental Section) are used for comparing gene rankings between predicted and ground-truth gene expression profiles.

Previous work^24^ formulates the gene expression prediction as a classification problem by classifying significantly regulated genes. Although such work showed promising results, the classification setting is actually not very effective and practical in the downstream applications because it cannot represent the whole gene expression profile. Subsequent analysis using chemical-induced gene expression profiles will benefit most from the information of the whole profiles. Thus, we target a more realistic but more challenging scenario in which the model predicts the ranking of genes in the gene expression profile. In particular, we evaluate CIGER, DeepCOP, and a random permutation (Supplementary Note 1) for the ranking task by measuring the ranking of up-regulated genes (genes that have z-scores *>* 0) and down-regulated genes (genes that have z-scores *<* 0). DeepCOP is not originally developed for predicting gene ranking so we use its predicted probability scores to generate ranked lists. As shown in Table 2, CIGER significantly outperforms DeepCOP and random permutation by a large margin across all ranking metrics. Specifically, CIGER achieves NDGC scores of 0.8275 and 0.8460 which reduces the error rates of DeepCOP by 9.1% and 6.8% for up-regulated and down-regulated gene ranking respectively. CIGER also achieves significantly better Precision@K compared to DeepCOP showing the effectiveness of CIGER for predicting the ranking of genes in the whole gene expression profiles of novel chemicals. To further validate the performances of CIGER, we conduct the significant testing (i.e., paired sample t-test) with respect to NDCG scores between CIGER and the best baseline method (i.e., DeepCOP). The p-values of the paired sample t-test for up-regulated and down-regulated gene ranking tasks are 1.93 × 10^−30^ and 4.18 × 10^−44^ respectively, thereby showing the superiority of CIGER for gene expression ranking compared to the existing methods. We also evaluate the performances of CIGER and baseline methods for ranking tasks with respect to each cell line. The cell-specific evaluations (i.e., NDCG and Precision@K) for these methods are shown in Supplementary Table S1 and S2.

**Table 2.** Average performances (NDCG and Precision@K) of CIGER, DeepCOP, and the random ranking for ranking up-regulated and down-regulated genes under the 5-fold cross-validation setting.

| Up-regulated gene ranking |  |  |  |  |  |
| --- | --- | --- | --- | --- | --- |
| Model | NDCG | P@10 | P@50 | P@100 | P@200 |
| Random | 0.7309 ± 0.0025 | 0.2045 ± 0.1270 | 0.2045 ± 0.0556 | 0.2045 ± 0.0382 | 0.2045 ± 0.0254 |
| TT-WOPT | 0.7384 ± 0.0010 | 0.2606 ± 0.0143 | 0.2395 ± 0.0093 | 0.2284 ± 0.0077 | 0.2181 ± 0.0060 |
| DeepCOP | 0.8083 ± 0.0022 | 0.5430 ± 0.0131 | 0.4656 ± 0.0111 | 0.4161 ± 0.0104 | 0.3559 ± 0.0068 |
| CIGER | **0.8275 ± 0.0041** | **0.5973 ± 0.0170** | **0.5276 ± 0.0126** | **0.4735 ± 0.0101** | **0.4027 ± 0.0077** |
| Down-regulated gene ranking |  |  |  |  |  |
| Model | NDCG | P@10 | P@50 | P@100 | P@200 |
| Random | 0.7418 ± 0.0013 | 0.2045 ± 0.1270 | 0.2045 ± 0.0556 | 0.2045 ± 0.0382 | 0.2045 ± 0.0254 |
| TT-WOPT | 0.7534 ± 0.0007 | 0.2876 ± 0.0076 | 0.2618 ± 0.0050 | 0.2471 ± 0.0052 | 0.2297 ± 0.0040 |
| DeepCOP | 0.8346 ± 0.0030 | 0.6084 ± 0.0190 | 0.5304 ± 0.0180 | 0.4779 ± 0.0143 | 0.4077 ± 0.0090 |
| CIGER | **0.8460 ± 0.0023** | **0.6342 ± 0.0120** | **0.5753 ± 0.0041** | **0.5250 ± 0.0034** | **0.4465 ± 0.0035** |

### Gene Expression Classification for *de novo* Chemicals

Besides the ranking setting, we also compare CIGER with baseline methods including TT-WOPT, DeepCOP, and logistic regression (LR) in the classification setting in which the models predict whether genes are up-regulated or down-regulated due to molecular intervention. As shown in Table 3, CIGER outperforms TT-WOPT, LR, and DeepCOP by a large margin which demonstrates its effectiveness for gene expression classification tasks. In particular, CIGER achieves AUC scores of 0.7202 and 0.7558 for up-regulated and down-regulated gene classification tasks respectively. For baseline methods, DeepCOP achieves better performances than LR indicating that the linear model is not capable to capture the relationship between input features and gene regulation effects. The performances of TT-WOPT for the two classification tasks, as we expected, are 0.4981 and 0.5096 which are equivalent to a coin toss. TT-WOPT designed for imputation setting does not leverage any feature information excepted the gene expression values in the training set when making predictions so this method is not suitable for *de novo* chemical setting. We also evaluate the classification performances of these models by AU-PRC and F1 scores. Supplementary Table S3 shows the results measured by these classification metrics.

**Table 3.** Average performances (AUC) of CIGER and baseline models for up-regulated and down-regulated gene classification tasks under 5-fold cross validation setting. (TT-WOPT, CIGER and its variants (i.e., CIGER^−A^ and CIGER^−NA^) are trained with z-score values while logistic regression and DeepCOP are trained with binary labels indicating gene regulation stages.)

| Methods | Objective Function | Classification Tasks |  |
| --- | --- | --- | --- |
|  |  | Up-regulated | Down-regulated |
| TT-WOPT | N/A | 0.4981 ± 0.0097 | 0.5096 ± 0.0097 |
| Logistic Regression | Binary Cross Entropy | 0.6270 ± 0.0209 | 0.6480 ± 0.0182 |
| DeepCOP | Binary Cross Entropy | 0.6764 ± 0.0176 | 0.6925 ± 0.0217 |
| CIGER<sup>-NA</sup> | ListMLE | 0.6741 ± 0.0071 | 0.7166 ± 0.0102 |
| CIGER<sup>-NA</sup> | ListNet | 0.6723 ± 0.0122 | 0.7254 ± 0.0081 |
| CIGER<sup>-NA</sup> | RankCosine | **0.6992 ± 0.0106** | **0.7289 ± 0.0123** |
| CIGER<sup>-NA</sup> | RankNet | 0.6810 ± 0.0052 | 0.7192 ± 0.0108 |
| CIGER<sup>-A</sup> | RankCosine | 0.7086 ± 0.0106 | 0.7448 ± 0.0040 |
| CIGER | RankCosine | **0.7202 ± 0.0057** | **0.7558 ± 0.0061** |

### Drug Repurposing for Pancreatic Cancer

#### Drug candidates prediction for pancreatic cancer

To further investigate the effectiveness of CIGER, we design the drug screening pipeline using this model to find potential treatment for pancreatic cancer from existing drugs. Previous drug repurposing researches have identified useful targets and drug candidates for killing pancreatic cancer cells or inhibiting tumor growths with limited numbers of drugs obtained from compound library or by target.^41,42^ Here we perform drug repurposing with all existing drugs from DrugBank. Furthermore, we aimed to discover drugs that can induce drug sensitivity of pancreatic cancer sub-types that are resistant to existing anti-cancer therapies rather than screen compounds that can kill cancer cells directly. A recent study has shown that the combination of metformin and vitamin C can restore TET2 and GATA6 activities in aggressive squamous-like pancreatic ductal adenocarcinoma sub-type, which are the biomarkers of classical-pancreatic tumor, thereby improving therapeutic responses and survival of aggressive pancreatic sub-type^43^. The main step of this screening pipeline is to compare the chemical-induced gene expression profiles generated by CIGER with gene expression profile computed from pancreatic cancer cell lines treated by metformin and vitamin C. For drug gene expression profiles, we send queries to the DrugBank database to retrieve the list of all existing drugs (i.e., 11179 drugs) with their corresponding SMILES representations and then use CIGER trained on the LINCS L1000 dataset to generate profiles for these drugs from their SMILES representations. For gene expression profile of pancreatic cancer treated by metformin and vitamin C, we perform differential expression analysis with DESeq2^44^ between metformin and vitamin C treated samples and mock-treated samples^43^. Then, we compute the similarity with respect to ranking information between the gene expression profiles of treatment and drugs across 10 cell lines by GSEA and Precision@200 scores to find potential treatment for this disease. Note that, we derive the treatment profile as the differential expression of treated disease samples versus untreated disease samples which is different from the differential expression of disease samples versus normal samples used in deriving the disease profile mentioned in previous works. Thus, the drug candidates from our pipeline would induce similar gene expression profiles as treatment (i.e., metformin/vitamin C) profile instead of having inverse correlation with the disease profile as in previous works. The details of the method used to generate drug and treatment gene expression profiles and the screening process are shown in Experimental Procedures section. Figure 3 shows the proposed drug screening pipeline using CIGER.

**Figure 3.**
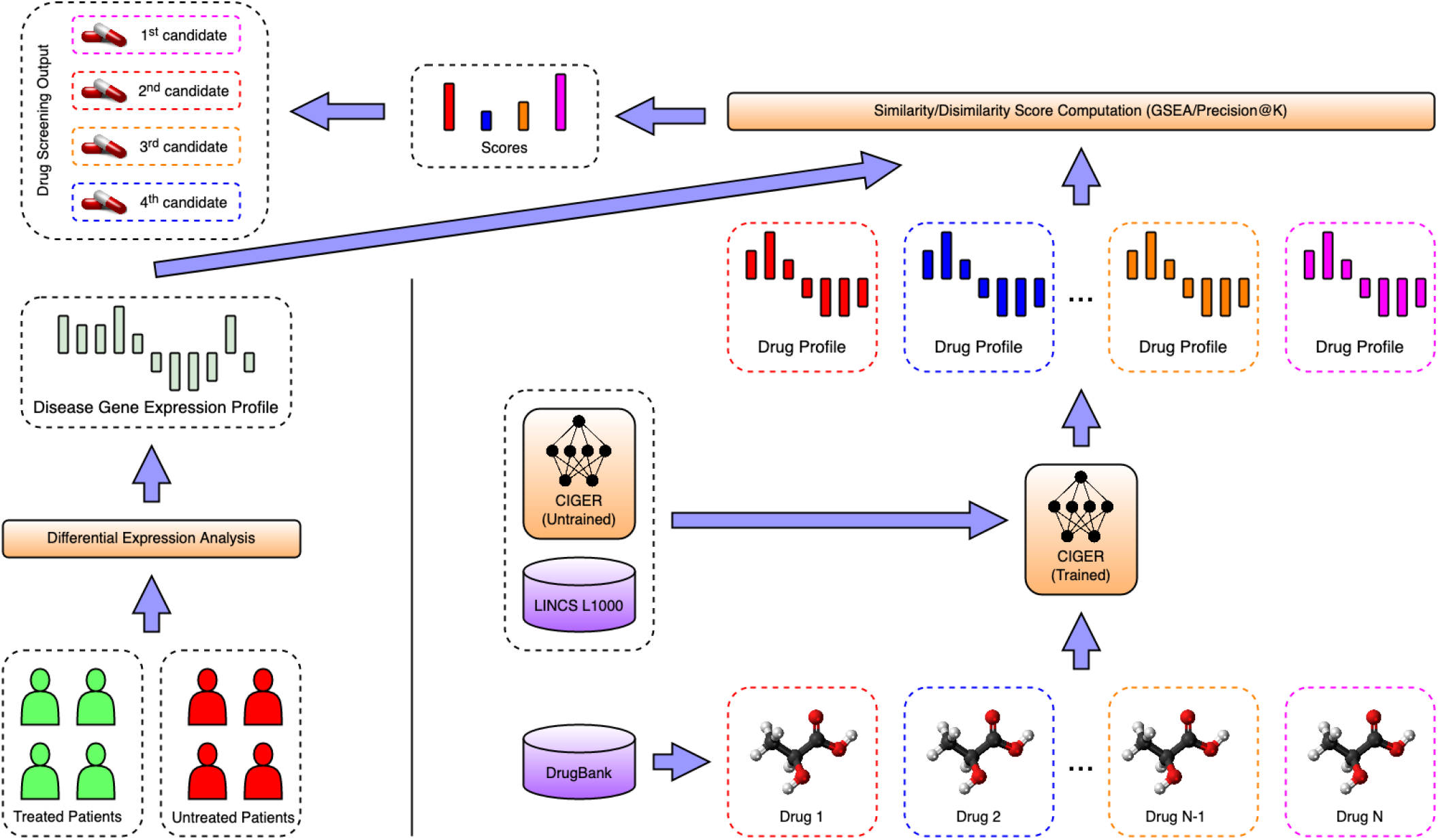
Drug screening pipeline using CIGER. This model is trained with the LINCS L1000 dataset to learn the relation between gene expression profiles and molecular structures (i.e., SMILES). Then molecular structures retrieved from the DrugBank database are put into CIGER to generate the corresponding gene expression profiles. Finally, these profiles are compared with treatment profiles calculated from treated and untreated samples to find the most potential treatments for that disease.

Top drugs selected by Precision@200 and GSEA scores are listed in Tables 4 and 5 respectively. The 2-dimensional molecular structures of these drugs are visualized in Supplementary Figures S3 (Precision@200) and S4 (GSEA). Sucrosofate and Inositol Hexasulphate in this list are known to bind human FGF1, which is related to tumor growth and invasion^45^. For drugs selected by GSEA score, 6 of them are confirmed to affect PI3K or mTOR pathway. As PI3K/AKT/mTOR signaling is one of the most important intracellular pathways that regulate the cell cycle, it can be targeted by drugs to regulate the metabolism in cancer cells, resulting in phenotype shift, increased cell death, and decreased cell proliferation^46,47^. Biguanide and its medication drug (i.e., Metformin) have been shown to be effective for pancreatic cancer tumor growth inhibition^43,48,49^. Note that the drugs selected by CIGER are not available in the training set of LINCS L1000 dataset, thereby showing its real potential application in drug discovery that enables high-throughput phenotypic drug screening by utilizing the molecular structure’s information only. Also, cell-specific prediction may also provide improvement for predictions. The cell-specific ranks and similarity scores (i.e., Precision@200 and GSEA) of these drug candidates are shown in Supplementary Tables S5 and S6.

**Table 4.** Drug candidates selected by Precision@200

| DrugBank ID | Name | Formula | Information |
| --- | --- | --- | --- |
| DB00364 | Sucralfate | $C_{12}H_{35}Al_9O_{55}S_8$ | Treat and prevent the return of duodenal ulcers |
| DB01666 | Inositol Hexasulphate | $C_6H_{12}O_{24}S_6$ | binding to human acidic fibroblast growth factor |
| DB14815 | Ginsenoside B2 | $C_{48}H_{82}O_{18}$ | extract from Panax notoginseng(Japanese ginseng), decreases the β-amyloid protein |
| DB15532 | Madecassoside | $C_{48}H_{78}O_{20}$ | found in Centella asiatica (Gotu kola), anti-inflammatory, wound healing, and anti-oxidant activities |
| DB06749 | Ginsenoside Rb1 | $C_{54}H_{92}O_{23}$ | abundant in Panax quinquefolius (American Ginseng), protected against amyloid β-induced neurotoxicity |
| DB14528 | Chromium gluconate | $C_{18}H_{33}CrO_{21}$ | supplement to intravenous solutions given for total parenteral nutrition |
| DB09517 | Sodium ferric gluconate complex | $C_{66}H_{121}Fe_2NaO_{65}$ | treat iron deficiency anemia |
| DB03995 | Betadex | $C_{42}H_{70}O_{35}$ | pharmacologically inactive substance, useful for stabilizing, solubilizing or delivering intermediate size molecules. |
| DB01901 | Sucrosofate | $C_{12}H_{22}O_{35}S_8$ | drug retention and drug encapsulation stability |

**Table 5.**
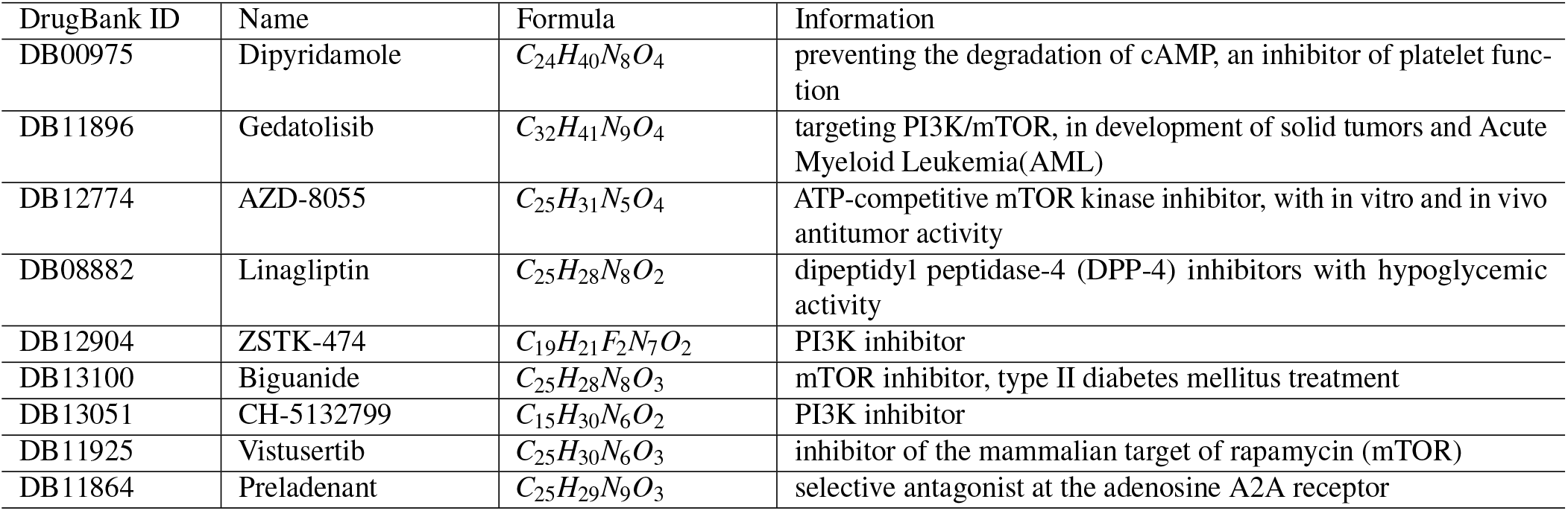
Drug candidates selected by GSEA

| DrugBank ID | Name | Formula | Information |
| --- | --- | --- | --- |
| DB00975 | Dipyridamole | $C_{24}H_{40}N_8O_4$ | preventing the degradation of cAMP, an inhibitor of platelet function |
| DB11896 | Gedatolisib | $C_{32}H_{41}N_9O_4$ | targeting PI3K/mTOR, in development of solid tumors and Acute Myeloid Leukemia(AML) |
| DB12774 | AZD-8055 | $C_{25}H_{31}N_5O_4$ | ATP-competitive mTOR kinase inhibitor, with in vitro and in vivo antitumor activity |
| DB08882 | Linagliptin | $C_{25}H_{28}N_8O_2$ | dipeptidyl peptidase-4 (DPP-4) inhibitors with hypoglycemic activity |
| DB12904 | ZSTK-474 | $C_{19}H_{21}F_2N_7O_2$ | PI3K inhibitor |
| DB13100 | Biguanide | $C_{25}H_{28}N_8O_3$ | mTOR inhibitor, type II diabetes mellitus treatment |
| DB13051 | CH-5132799 | $C_{15}H_{30}N_6O_2$ | PI3K inhibitor |
| DB11925 | Vistusertib | $C_{25}H_{30}N_6O_3$ | inhibitor of the mammalian target of rapamycin (mTOR) |
| DB11864 | Preladenant | $C_{25}H_{29}N_9O_3$ | selective antagonist at the adenosine A2A receptor |

#### Experimental validation for pancreatic cancer drug candidates

To evaluate our candidates generated from our predictions, several drugs including Dipyridamole, AZD-8055, Linagliptin, Preladenant are tested *in vitro* together with the combination of Metformin and Vitamin C as a positive control. We used the above drugs to treat pancreatic cancer cell lines and perform western blot to show the level of GATA6 and TET2, thus evaluating the effect of the predicted candidate drugs.

As shown in Figures 4A and 4B, western blot following quantification showed that the combination of Metformin and Vitamin C increased TET2 and GAT6 levels in PANC-1 cells at 24 hours. Dipyridamole can also significantly increase TET2 level after 24 hours treatments in PANC-1 cells, and Linagliptin increased both TET2 and GATA6 levels significantly, suggesting that they can induce similar response to Metformin and Vitamin C.

**Figure 4.**
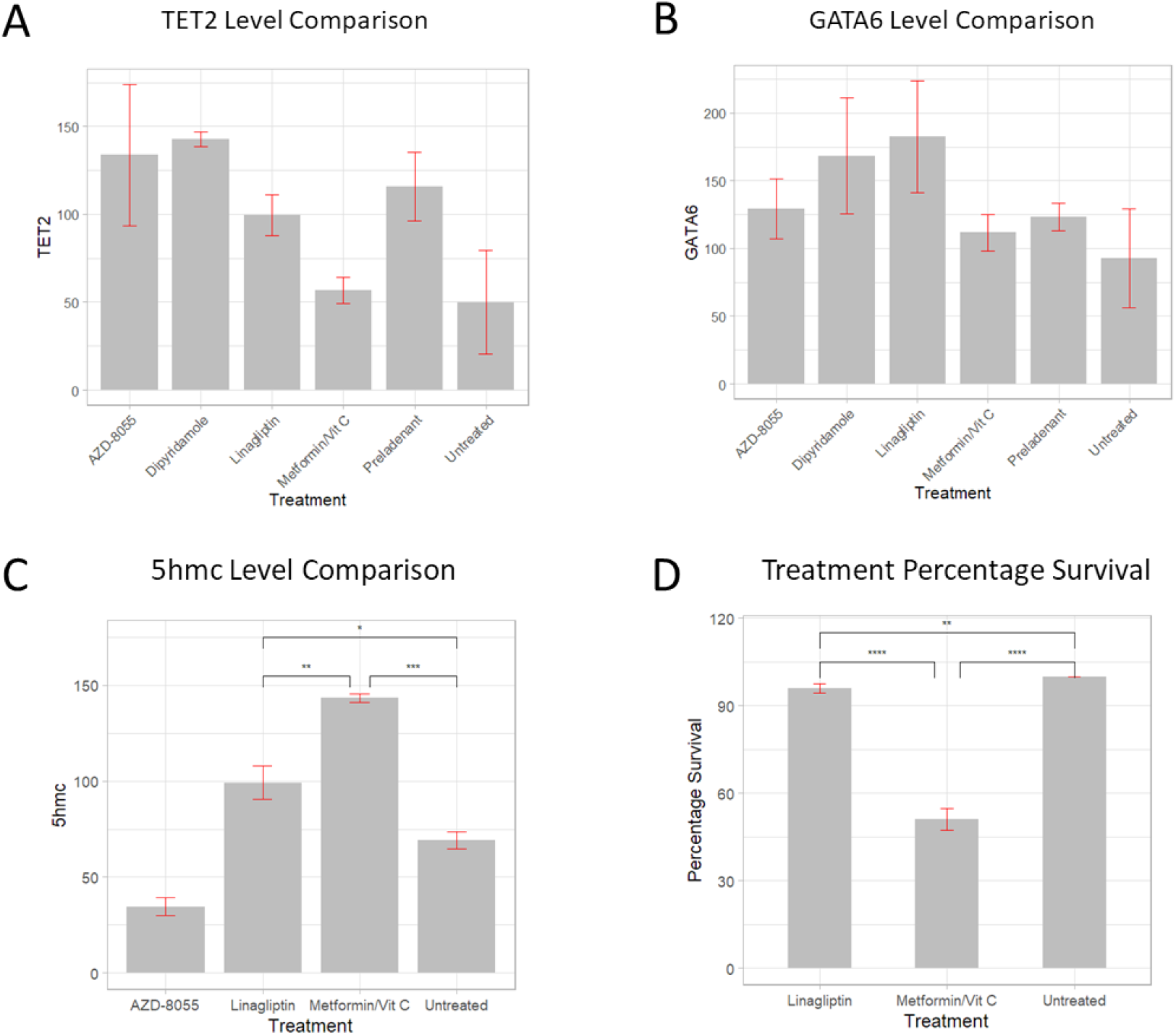
(A) Quantification of TET2 levels in drug treatments. Dipyridamole and Linagliptin can significantly increase TET2 level after 24 hours treatments. (B) Quantifications of GATA6 expressions in drugs treatment. Linagliptin increased GATA6 expressions in PANC-1 after 24hr treatment. Data are presented as mean ± SD (n=3). (C) Linagliptin and Metformin Vitamin C increased 5hmc levels in PANC-1 cells after 24-hour treatment. Quantifications of 5hmc dot blots (n=3), data are represented as mean± SD. *p<0.05, **p<0.01, ***p<0.001 (Unpaired two-tailed t test and one-way ANOVA) (D) The effect of drug treatments on clonogenic survival as a measure of growth rate. All data are presented as mean ± SD (n=3). **p<0.005, ****p<0.0001 (analysed using one-way ANOVA)

To investigate whether the increase in TET2 and GATA6 would have an effect on 5hmc, dot plots of all drugs were performed and quantified. As shown in Figure 4C, Metformin/Vitamin C and Linagliptin significantly increased 5hmc level in PANC-1 genomic DNA after 24-hour treatment. To study the effect of treatments on the growth of PANC-1 cells, clonogenic survival of cells with Metformin/Vitamin C or Linagliptin treatment was analyzed. As showed in Figure 4D, Metformin and Vitamin C treatment demonstrated a significantly lower percentage of survival compared to negative control, and Linagliptin treatment caused a significantly lower survival rate compared to both the negative control and the combination of metformin and Vitamin C. To understand the way Linagliptin improve treatment sensitivity, we analysis the predicted drug signature of Linagliptin with paslincs^50^ to find the affected pathways. We found antifolate resistance pathway on top of the list, which is related to drug resistance in cancer treatment. The compounds we identified can be further studied *in vivo*, or use tools like Code-AE^51^ to predict their patient-specific clinical response to predict their performance in the real application.

### Ablation Study for CIGER

An ablation study is conducted to further investigate how CIGER surpasses the limitations of existing methods for chemical-induced gene expression prediction. In particular, we remove components from CIGER (i.e., CIGER^−NA^ and CIGER^−A^) and observe the changes in its prediction performances. We also explore the impact of noisy data on prediction performance.

#### Learning-to-Rank Objective Function

In this experiment, we investigate the effectiveness of learning-to-rank objective functions (i.e., ListMLE^32^, ListNet^31^, RankCosine^33^, and RankNet^28^) for learning the dependencies among genes. In order to do that, we compare CIGER^−NA^ with DeepCOP (Binary Cross Entropy) for gene expression classification tasks. CIGER^−NA^ is a variant of CIGER in which the attention component is removed and extended connectivity fingerprint (ECFP) is used instead of neural fingerprint (learned by graph convolutional network) to represent chemical so the main difference between CIGER^−NA^ and DeepCOP is at the objective functions they optimize. As shown in Table 3, the overall performance of CIGER^−NA^ is better than DeepCOP for both of the two classification tasks. Among these objective functions, using CIGER^−NA^ with RankCosine achieves the best improvement compared to DeepCOP. In particular, it achieves AUC scores of 0.6992 and 0.7289 which is significantly better than AUC scores of 0.6764 and 0.6925 of DeepCOP for up-regulated and down-regulated gene classification tasks respectively. Therefore, we use RankCosine as the ranking objective function for CIGER and its variant.

#### Data-driven Representations for Chemicals

To validate the improvement of data-driven features over pre-defined features for chemicals, we compare the performances of using ECFP (i.e., CIGER^−NA^) and neural fingerprint generated by graph convolutional network (i.e., CIGER^−A^). As shown in Table 3, using neural fingerprint achieves better performances than ECFP. In particular, CIGER^−A^ achieves AUC scores of 0.7086 and 0.7448, which are better compared to 0.6992 and 0.7289 of CIGER^−NA^ indicating the effectiveness of the approach that automatically learns representations for chemicals from data.

#### Multi-head Attention Mechanism

We compare CIGER with its variants in this experiment to evaluate the effectiveness of the multi-head attention component for prediction performance. As shown in Table 3, by leveraging the attention mechanism to learn the dependencies among genes and chemicals, CIGER achieves the best performances compared to its variants. In particular, it outperforms CIGER^−A^ by achieving AUC scores of 0.7202 and 0.7558 compared to 0.7086 and 0.7448 of CIGER^−A^ for up-regulated and down-regulated gene classification tasks, respectively. CIGER^−NA^, without both graph convolutional network and multi-head attention components, as we expected, achieve the worst performances among its variants. All these results demonstrate the improvement of using multi-head attention for gene expression prediction.

#### Noisy Gene Expression Data

To validate the impact of the noisy problem in LINCS L1000 dataset on the prediction performance of CIGER, we train this model on the whole dataset (i.e., without removing noisy profiles which have APC scores < 0.6 among their bio-replicates) and compare with the one trained on high-quality data only (i.e., including only profiles which have APC scores > 0.6 among their bio-replicates). The results (i.e., NDCG, Precision@K) shown in Supplementary Table S4 indicates that noisy gene expression can significantly hinder the prediction performance of CIGER. In particular, its NDCG scores decrease from 0.8275 and 0.8460 to 0.7761 and 0.7966 for up-regulated and down-regulated gene ranking respectively.

### Existing Limitations

Although achieving superior results compared to the baseline methods and showing feasibility in gene expression-based drug repurposing for pancreatic cancer, our proposed method still has some limitations. First, it cannot generate chemical-induced gene expression profiles with respect to the new cell lines (i.e., except 10 cell lines in the training dataset) caused by the lack of cell line representations. Second, due to the noisy issue in LINCS L1000 dataset, we can only utilize a small subset of this data for training thereby hindering the prediction performance of our method for *de novo* chemicals. Third, the limited size of LINCS L1000 dataset in terms of chemicals (i.e., ~ 20, 000 small molecules) inhibits CIGER to learn generalized representation for chemicals in the complex molecular space. Finding efficient cell line representation, denoising gene expression data, and pre-training on the large molecular datasets are the keys to surpassing these limitations and we leave them in our future works.

## Conclusion

Large-scale analysis that leverages chemical-induced gene expression profiles has attracted great attention in drug discovery. However, the effectiveness of this approach is limited by the spareness and noisy measurement problems. In this study, we propose CIGER - a novel and robust neural network-based model for predicting the ranking of genes in the gene expression profiles induced by *de novo* chemicals. Our model achieves state-of-the-art results compared to other methods for both gene expression classification and ranking tasks in *de novo* chemical setting. Furthermore, with the capability of predicting the ranking of genes in the chemical-induced gene expression profiles across different cell lines leveraging the chemical structures only, CIGER provides new opportunities on subsequent molecular phenotype-based drug repurposing by comparing the ranking of genes in chemical-induced profiles with the treatment profiles computed from chemical-treated and untreated disease states. The similar or reverse of gene expression ranking will suggest the most potential drug candidates for specific diseases. In summary, CIGER could be a powerful tool for phenotypic compound screening.

## Experimental Procedures

### Ranking task definition

In LINCS L1000 dataset, each experiment can be considered as a tuple of chemical compound, cell line, and the corresponding gene expression profile. These biological and chemical objects are transformed to numerical representations for using in the computational models. In particular, the L1000 dataset can be represented by the following matrices 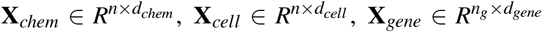, and 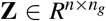 where **X**_*chem*_, **X**_*cell*_, **X**_*gene*_ are feature matrices of chemicals, cell lines, and L1000 genes in the dataset, 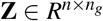 is the gene expression matrix, *n, n*_*g*_ are numbers of experiments and L1000 genes, and *d*_*chem*_, *d*_*cell*_, *d*_*gene*_ are feature dimensions of chemical compound, cell line, and gene respectively. The goal of this task is predicting the ranking of genes in the expression profile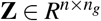 based on the feature matrices.

### CIGER architecture

This consists of four main components: (1) the feature mapping component that transforms biological and chemical objects to numerical representations including graph convolutional network to transform simplified molecular-input line-entry system (SMILES) representations of chemicals to numerical vectors and embedding lookup tables to transform cell lines and L1000 gene indexes to binary vectors, (2) the attention component that looks at all L1000 genes and chemical to create contextualized representation for each gene, (3) the prediction component that predicts the ranking of all L1000 gene in gene expression, and (4) the learning-to-rank objective function that optimizes the global prediction performance for all L1000 genes. Figure 1 presents the overview architecture of CIGER. The details of each component are as follows.

#### Feature Mapping for Biological and Chemical Objects

We use graph convolutional network and gene ontology consortium to construct numerical representations for chemical compounds and L1000 genes. For cell lines, we simply use onehot vectors.

The chemical feature matrix **X**_*chem*_ is generally pre-defined in traditional approaches. One popular method is extended connectivity fingerprint (ECFP) that represents molecular substructures by means of circular atom neighborhoods. Specifically, the presence or absence of sub-structures is encoded in a fixed-size binary vector. The main drawback of this method is the sub-structures need to be available before training, and therefore, may not be the optimized way to represent the chemicals for particular tasks. Recently, with the advancement of graph neural networks, some data-driven methods have been proposed to effectively exploit the graph-based structure of chemical^52,53^. Compared to pre-defined approaches, these methods can automatically find the most important sub-structures which are optimized representations for chemicals for the prediction tasks by optimizing the objective function from training. In our work, we use graph convolutional network^35^ to exploit information from chemicals which can be seen as graphs of atoms (nodes) and bonds (edges). This method can be seen as the differentiable variant of ECFP in which every step is continuous and differentiable, and therefore, allows updates from gradient propagation. In particular, graph convolutional network updates the representation of one particular node from the information of its neighborhoods in the graph by convolutional operation so each node in the output layer represents the sub-structure of the original graph. Following the setting in^35^, we use the 2-layer graph convolutional network (radius = 2) which means that the sub-structures represented by this method are the span of 2-hop distance from the atom. Inputs for graph convolutional network are the feature vectors of atoms and bonds that captures their properties such as atom symbol, degree, and type of bonds. The dimension of fingerprints generated by graph convolutional network is set to be 1024 which is similar to ECFP for a fair comparison. The detailed implementation of the graph convolution network used for chemicals are shown in Supplementary Note 2.

Gene ontology consortium^54^ has been shown to be an effective way to represent genes and proteins by capturing their biological process, molecular function, and cellular component. In our experiments, we follow the data processing described in^24^ by selecting 1107 gene ontology terms that appeared in at least three L1000 genes and using them to construct **X**_*gene*_. These representations can be seen as binary vectors where the indexes of bit 1 mean the appearance of gene ontology terms associated with these indexes.

#### Multi-head Attention for Contextualized Representation

We utilize multi-head attention^34^ to capture the dependencies among genes for learning better representation. In particular, for *i-th* experiment, chemical 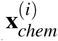 and cell line 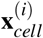 are concatenated with each gene 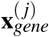 in **X***gene* and then putted into feed-forward neural network layer and ReLU activation function to generate contextualized 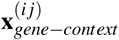 as follows:

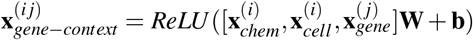

where 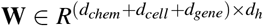, **b** ∈ *d*_*h*_ are learned parameters and [*a, b*] is concatenation operation on *a, b*. The contextualized representation for L1000 genes are packed into matrix 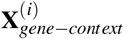 and then are putted into multi-head attention to learn attention-based representation. In particular, multi-head attention transforms the input feature matrix (i.e. 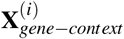) to three separate matrices 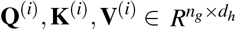 as follows:

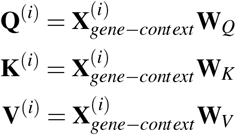

Where 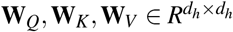 are the trainable parameter matrices, *d*_*h*_ is dimension of transformed features and then the attention-based representations for the input features are computed as follows:

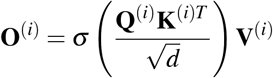

Where 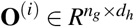, *σ*, and *d* are the attention representation, softmax function, and scale factor.

#### Multi-output Prediction for Gene Ranking

The 2-layer feed-forward neural network with ReLU activation function is used to predict the rank of each gene in the gene expression profile. The weight of this network is shared across all L1000 genes.

#### Learning-to-rank Objective Functions

CIGER optimizes the predictions for all L1000 genes together rather than individually by using learning-to-rank objective functions. In particular, CIGER treats the gene expression profiles as the lists ranked by their z-score and then minimizes several learning-to-rank objective functions including both pair-wise (i.e. RankNet^28^) and list-wise (i.e. ListNet^31^, ListMLE^32^, and RankCosine^33^) functions between the predicted (**y**) and the ground-truth (**z**) gene expression profiles. Details of these objective functions are presented in Supplementary Note 3.

### Baseline methods

We compare CIGER with the following baseline models for chemical-induced gene expression ranking and classification tasks.

#### Logistic Regression (LR)

The linear model used in gene expression classification task. We use the scikit-learn implementation^55^ to train this model on LINCS L1000 dataset. Inputs for LR are the concatenations of 1024-bit circular topological fingerprints for chemical, one-hot vector for cell line, and multi-hot vectors (i.e., 1107-bit) that represent the inclusion of Gene Ontology terms for L1000 genes. The outputs of linear functions are put into the logistic function to model the probabilities of being (up or down) regulated for L1000 genes induced by chemicals.

#### DeepCOP^24^

The neural network-based model for gene expression classification task. We reimplement this model in PyTorch framework^56^ and use the same hyper-parameters as in the original paper. This model consists of three layers with SeLU activation function for the first layer and ReLU activation function for the following layers. Inputs for DeepCOP are similar to LR and the objective function is binary cross-entropy between the ground-truth and predicted labels.

#### Tensor-train Weight Optimization (TT-WOPT)^40^

The tensor completion-based model used to impute missing values in high-dimensional (tensor) data from existing values. It has shown good performance when applying to predict z-score values of the LINCS L1000 dataset. This method leverages existing labels (z-score) only to make predictions so additional feature information such as chemical, cell lines, and genes are not required. We use the MatLab implementation provided by the authors to train this model in our *de novo* chemical setting.

#### CIGER^−A^

The variant of our proposed model that makes predictions without attention mechanism.

#### CIGER^−NA^

The variant of our proposed model that does not use both neural fingerprint and attention mechanism.

### Evaluation metrics

To evaluate the performances of prediction models on the testing sets, the area under the receiver operating characteristic (AUC) is chosen for classification tasks, and NDCG and Precision@K are used for ranking tasks. The details of NDCG and Precision@K are as follows.

#### Normalized Discounted Cumulative Gain (NDCG)

is the metric used to evaluate the performances of models in ranking tasks. This metric focuses on two aspects of the ranking models including (1) giving higher ranks for higher relevant items and (2) highly relevant items are ranked higher than marginally relevant items, and in turn, having higher ranks than non-relevant items. In particular, NDCG at rank p is calculated as follows:

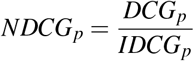

where Discounted Cumulative Gain (*DCG*_*p*_) and Ideal Discounted Cumulative Gain (*IDCG*_*p*_) which is max possible value of *DCG*_*p*_ are computed as follows:

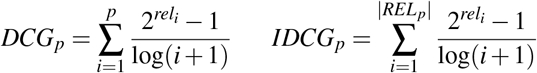

where *rel*_*i*_ is the relevant score of the result at position i and *REL*_*p*_ is the sorted list of relevant items up to position p. In our setting, the relevant scores are z-score (minus z-score in the case of ranking down-regulated genes) values in the gene expression profiles. Because negative scores cause NDCG to be unbounded so for up-regulated and down-regulated gene rankings, we set all negative relevant score to be 0.

#### Precision@K

is another metric we used to evaluate the performances of models in ranking tasks. It is the proportion of genes in the top-k predicted set that is up-regulated or down-regulated. In particular, Precision@K is computed as follows:

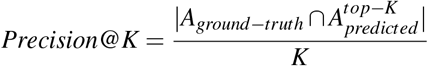

where *A*_*ground*−*truth*_ is the set of up-regulated or down-regulated genes (i.e., we select the top 200 genes that have the largest and smallest z-scores as the sets of the ground-truth up-regulated and down-regulated genes respectively) and 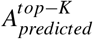 is the sets of top-K genes in the predicted ranked lists. In our study, we evaluate the performances of models at different K-levels including 10, 50, 100, and 200.

### Drug Repurposing Pipeline

#### Drug Gene Expression Profile

To generate drug profiles used in the drug screening process, we send queries to the DrugBank database to retrieve the list of all existing drugs (i.e., 11179 drugs) with their corresponding SMILES representations and then use CIGER trained on the LINCS L1000 dataset to generate gene expression profiles (i.e., gene ranking) for these drugs from their SMILES representations. In our study, each drug is represented by 10 cell-specific gene expression profiles. Note that under cross-validation setting, each profile is the average of the corresponding profiles generated from different models trained on different data folds. In summary, this process results in a ranking matrix of 11179 rows (drugs) and 10 columns (cell lines).

#### Treatment Gene Expression Profile

For pancreatic cancer treatment gene expression profile, we perform differential expression analysis between metformin and vitamin C treated samples and mock-treated samples with DESeq2^44^. A recent study showed that metformin and vitamin C treatment can restore TET2 activity in aggressive squamous-like pancreatic ductal adenocarcinoma subtype, and increase biomarker of classical-pancreatic tumor, which are related to improved therapeutic responses and survival^43^. To search for drugs that can restore epigenetic control in pancreatic tumors like metformin and vitamin C, we used pancreatic cancer gene expression data from this study, where the human pancreatic tumor cell line PSN1, orthotopically implanted into mice, was treated with Metformin combined with vitamin C or mock treatment. RNA-seq gene expression data are filtered to get human-only reads. In total, 3 Metformin/vitamin C treated samples and 3 mock-treated control samples are used for differential expression analysis. Those up/down-regulated genes can be used as signatures that characterize the treatment (i.e., the differential expression analysis is from mock-treated samples to treated samples).

#### Screening Method

A screening process is conducted by comparing drug profiles with treatment profile in terms of gene ranking. Specifically, Precision@200 and GSEA scores are used to find drugs whose profiles are most similar to the treatment profiles with respect to each cell line. The reason is that we want to find drugs that induce similar responses in pancreatic cancer as metformin/vitamin C treatment which has been shown to be effective in restoring epigenetic control in pancreatic cancer cell and improving therapeutic responses. Then, for each cell line, the top 10 most similar drug candidates are retrieved for further analysis. Previous studies showed that consensus gene expression profiles can give a more comprehensive representation and improve confidence in gene-expression analysis so we use results from all 10 cell lines to determine drug candidates. In particular, we select drugs that are in the top 10 of at least 3 cell lines for Precision@200 and 2 cell lines for GSEA as our potential treatments for pancreatic cancer.

## References

1. Terstappen, G. C., Schlüpen, C., Raggiaschi, R. & Gaviraghi, G. Target deconvolution strategies in drug discovery. Nat. Rev. Drug Discov. 6, 891–903 (2007).

2. Lamb, J. et al. The connectivity map: using gene-expression signatures to connect small molecules, genes, and disease. Science 313, 1929–1935 (2006).

3. Hu, G. & Agarwal, P. Human disease-drug network based on genomic expression profiles. PLOS One 4 (2009).

4. Dudley, J. T., Deshpande, T. & Butte, A. J. Exploiting drug–disease relationships for computational drug repositioning. Briefings Bioinforma. 12, 303–311 (2011).

5. Kosaka, T. et al. Identification of drug candidate against prostate cancer from the aspect of somatic cell reprogramming. Cancer Sci. 104, 1017–1026 (2013).

6. Wei, G. et al. Gene expression-based chemical genomics identifies rapamycin as a modulator of mcl1 and glucocorticoid resistance. Cancer Cell 10, 331–342 (2006).

7. Hassane, D. C. et al. Discovery of agents that eradicate leukemia stem cells using an in silico screen of public gene expression data. Blood 111, 5654–5662 (2008).

8. Stegmaier, K. et al. Gene expression–based high-throughput screening (ge-hts) and application to leukemia differentiation. Nat. Genet. 36, 257–263 (2004).

9. Lamb, J. The connectivity map: a new tool for biomedical research. Nat. Rev. Cancer 7, 54–60 (2007).

10. Chong, C. R. & Sullivan, D. J. New uses for old drugs. Nature 448, 645–646 (2007).

11. Novac, N. Challenges and opportunities of drug repositioning. Trends Pharmacol. Sci. 34, 267–272 (2013).

12. Subramanian, A. et al. A next generation connectivity map: L1000 platform and the first 1,000,000 profiles. Cell 171, 1437–1452 (2017).

13. Hodos, R. et al. Cell-specific prediction and application of drug-induced gene expression profiles. In Pacific Symposium on Biocomputing, vol. 23, 32–43 (World Scientific, 2018).

14. Yue, X. et al. Graph embedding on biomedical networks: methods, applications and evaluations. Bioinformatics 36, 1241–1251 (2020).

15. Troyanskaya, O. et al. Missing value estimation methods for dna microarrays. Bioinformatics 17, 520–525 (2001).

16. Bø, T. H., Dysvik, B. & Jonassen, I. Lsimpute: accurate estimation of missing values in microarray data with least squares methods. Nucleic Acids Res. 32, e34–e34 (2004).

17. Kim, H., Golub, G. H. & Park, H. Missing value estimation for dna microarray gene expression data: local least squares imputation. Bioinformatics 21, 187–198 (2005).

18. Cai, Z., Heydari, M. & Lin, G. Iterated local least squares microarray missing value imputation. J. Bioinforma. Comput. Biol. 4, 935–957 (2006).

19. Oba, S. et al. A bayesian missing value estimation method for gene expression profile data. Bioinformatics 19, 2088–2096 (2003).

20. Wang, X., Li, A., Jiang, Z. & Feng, H. Missing value estimation for dna microarray gene expression data by support vector regression imputation and orthogonal coding scheme. BMC Bioinforma. 7, 32 (2006).

21. Ouyang, M., Welsh, W. J. & Georgopoulos, P. Gaussian mixture clustering and imputation of microarray data. Bioinformatics 20, 917–923 (2004).

22. Lagunin, A., Ivanov, S., Rudik, A., Filimonov, D. & Poroikov, V. Digep-pred: web service for in silico prediction of drug-induced gene expression profiles based on structural formula. Bioinformatics 29, 2062–2063 (2013).

23. Iwata, M., Sawada, R., Iwata, H., Kotera, M. & Yamanishi, Y. Elucidating the modes of action for bioactive compounds in a cell-specific manner by large-scale chemically-induced transcriptomics. Sci. Reports 7, 40164 (2017).

24. Woo, G. et al. Deepcop: deep learning-based approach to predict gene regulating effects of small molecules. Bioinformatics 36, 813–818 (2020).

25. Bourdakou, M. M., Athanasiadis, E. I. & Spyrou, G. M. Discovering gene re-ranking efficiency and conserved gene-gene relationships derived from gene co-expression network analysis on breast cancer data. Sci. Reports 6, 1–29 (2016).

26. Rendle, S., Freudenthaler, C., Gantner, Z. & Schmidt-Thieme, L. Bpr: Bayesian personalized ranking from implicit feedback. In Proceedings of the Twenty-Fifth Conference on Uncertainty in Artificial Intelligence, 452–461 (2009).

27. Wang, Y., Sun, H. & Zhang, R. Adamf: Adaptive boosting matrix factorization for recommender system. In International Conference on Web-Age Information Management, 43–54 (Springer, 2014).

28. Burges, C. et al. Learning to rank using gradient descent. In Proceedings of the 22nd International Conference on Machine Learning, 89–96 (2005).

29. Freund, Y., Iyer, R., Schapire, R. E. & Singer, Y. An efficient boosting algorithm for combining preferences. J. Mach. Learn. Res. 4, 933–969 (2003).

30. Cao, Y. et al. Adapting ranking svm to document retrieval. In Proceedings of the 29th Annual International ACM SIGIR Conference on Research and Development in Information Retrieval, 186–193 (2006).

31. Cao, Z., Qin, T., Liu, T.-Y., Tsai, M.-F. & Li, H. Learning to rank: from pairwise approach to listwise approach. In Proceedings of the 24th International Conference on Machine Learning, 129–136 (2007).

32. Xia, F., Liu, T.-Y., Wang, J., Zhang, W. & Li, H. Listwise approach to learning to rank: theory and algorithm. In Proceedings of the 25th International Conference on Machine Learning, 1192–1199 (2008).

33. Qin, T. et al. Query-level loss functions for information retrieval. Inf. Process. & Manag. 44, 838–855 (2008).

34. Vaswani, A. et al. Attention is all you need. In Advances in Neural Information Processing Systems, 5998–6008 (2017).

35. Duvenaud, D. K. et al. Convolutional networks on graphs for learning molecular fingerprints. In Advances in Neural Information Processing Systems, 2224–2232 (2015).

36. Liu, C. et al. Compound signature detection on lincs l1000 big data. Mol. BioSystems 11, 714–722 (2015).

37. Li, Z., Li, J. & Yu, P. l1kdeconv: an r package for peak calling analysis with lincs l1000 data. BMC Bioinforma. 18, 356 (2017).

38. Duan, Q. et al. L1000cds 2: Lincs l1000 characteristic direction signatures search engine. NPJ Syst. Biol. Appl. 2, 1–12 (2016).

39. Qiu, Y., Lu, T., Lim, H. & Xie, L. A Bayesian approach to accurate and robust signature detection on LINCS L1000 data. Bioinformatics (2020).

40. Iwata, M. et al. Predicting drug-induced transcriptome responses of a wide range of human cell lines by a novel tensor-train decomposition algorithm. Bioinformatics 35, i191–i199 (2019).

41. Fujii, A. et al. The novel driver gene asap2 is a potential druggable target in pancreatic cancer. Cancer Sci. 112, 1655 (2021).

42. Kim, I. et al. A drug-repositioning screen for primary pancreatic ductal adenocarcinoma cells identifies 6-thioguanine as an effective therapeutic agent for tpmt-low cancer cells. Mol. Oncol. 12, 1526–1539 (2018).

43. Eyres, M. et al. Tet2 drives 5hmc marking of gata6 and epigenetically defines pancreatic ductal adenocarcinoma transcriptional subtypes. Gastroenterology (2021).

44. Love, M. I., Huber, W. & Anders, S. Moderated estimation of fold change and dispersion for rna-seq data with deseq2. Genome Biol. 15, 550 (2014).

45. Bai, Y.-P. et al. Fgf-1/-3/fgfr 4 signaling in cancer-associated fibroblasts promotes tumor progression in colon cancer through erk and mmp-7. Cancer Sci. 106, 1278–1287 (2015).

46. Xie, Y. et al. Phosphorylation of gata-6 is required for vascular smooth muscle cell differentiation after mtorc1 inhibition. Sci. Signal. 8, ra44–ra44 (2015).

47. Liu, R., Leslie, K. L. & Martin, K. A. Epigenetic regulation of smooth muscle cell plasticity. Biochimica et Biophys. Acta (BBA)-Gene Regul. Mech. 1849, 448–453 (2015).

48. Li, X., Li, T., Liu, Z., Gou, S. & Wang, C. The effect of metformin on survival of patients with pancreatic cancer: a meta-analysis. Sci. Reports 7, 1–8 (2017).

49. Hébert, A. et al. Phenylethynylbenzyl-modified biguanides inhibit pancreatic cancer tumor growth. Sci. Reports 11, 1–11 (2021).

50. Ren, Y. et al. Predicting mechanism of action of cellular perturbations with pathway activity signatures. Bioinformatics 36, 4781–4788 (2020).

51. He, D., Liu, Q. & Xie, L. Robust prediction of patientspecific clinical response to unseen drugs from in vitro screens using context-aware deconfounding autoencoder. bioRxiv (2021).

52. Gilmer, J., Schoenholz, S. S., Riley, P. F., Vinyals, O. & Dahl, G. E. Neural message passing for quantum chemistry. In Proceedings of the 34th International Conference on Machine Learning, 1263–1272 (PMLR, 2017).

53. Yang, K. et al. Analyzing learned molecular representations for property prediction. J. Chem. Inf. Model. 59, 3370–3388 (2019).

54. Ashburner, M. et al. Gene ontology: tool for the unification of biology. Nat. Genet. 25, 25–29 (2000).

55. Pedregosa, F. et al. Scikit-learn: Machine learning in Python. J. Mach. Learn. Res. 12, 2825–2830 (2011).

56. Paszke, A. et al. Automatic differentiation in pytorch. In Proceedings of the 2017 Neural Information Processing Systems Workshop Autodiff (2017).

