## Supplementary Information for "Chemical-induced Gene Expression Ranking and its Application to Pancreatic Cancer Drug Repurposing"

[1]Department of Computer Science and Engineering, The Ohio State University, Columbus, 43210, USA
[2]Ph.D. Program in Biology, The Graduate Center, The City University of New York, New York, 10016, USA
[3]Department of Oncology, University of Oxford, Oxford, OX3 7DQ, UK
[4]Epicombi Therapeutics, Charlbury, OX7 3SB, UK
[5]Department of Computer Science, Hunter College, The City University of New York, New York, 10065, USA
[6]Ph.D. Program in Computer Science and Biochemistry, The Graduate Center, The City University of New York, New York, 10016, USA
[7]Helen and Robert Appel Alzheimer's Disease Research Institute, Feil Family Brain & Mind Research Institute, Weill Cornell Medicine, Cornell University, New York, 10021, USA
[8]Department of Biomedical Informatics, The Ohio State University, Columbus, 43210, USA
[9]Translational Data Analytics institute, The Ohio State University, Columbus, 43210, USA

### ABSTRACT

This supplementary document includes supplementary notes, figures, and tables that support the manuscript "Chemical-induced Gene Expression Ranking and its Application to Pancreatic Cancer Drug Repurposing".

### Supplementary Notes

#### Supplementary Note 1: Random Ranking Method

The random ranking method can be seen as the random permutation over the gene sequence. Thus, its performances measured by Precision@K can be computed theoretically. In particular, Precision@K returned by the random ranking method can be seen as the random variable $X = Y/K$ where $Y$ follows the hypergeometric distribution $f(y|A,N,n)$ with $A = K$ is the number of top-K regulated genes, $N = n_g$ is the number of genes, and $n = K$ is number of top regulated genes in predicted profiles used to compute Precision@K. Then the average performance of the random ranking method measured by Precision@K is computed as $\bar{X} = \frac{\bar{Y}}{K} = \frac{nA}{NK} = \frac{K}{n_g}$. For measuring performances by NDCG we run random ranking method 100 times for each gene expression profile and return the average results.

#### Supplementary Note 2: Details of Data-driven Graph-based Fingerprint (Neural Fingerprint)

The pseudo-code of data-driven graph-based fingerprint generated by graph convolutional network is shown in Algorithm 1. In general, graph convolutional network updates the representation of one particular nodes from information of its neighborhoods in the graph by convolutional operation so each node in the output layer represents the sub-structure of the original graph. Follow the setting in[1], we use the 2-layer graph convolutional network (radius = 2) which means that the sub-structures represented by this method are the span of 2-hop distance from the atom. Inputs for graph convolutional network are the feature vectors of atoms and bonds that captures their properties such as atom symbol, degree, and type of bonds. The dimension of fingerprints generated by graph convolutional network is set to be 1024 which is similar to ECFP for a fair comparison.

**Algorithm 1:** Pseudo-code of data-driven graph-based fingerprint

**Input:** Chemical graph = $(V,E)$, radius $R$, hidden weights $(\mathbf{H}_1^1,...,\mathbf{H}_R^5),(\mathbf{U}_1,...,\mathbf{U}_l),(\mathbf{W}_1,...,\mathbf{W}_l)$
**Output:** Neural fingerprint $\mathbf{f}$
**for** $l = 1$ *to* $R$ **do**
  **for** $i = 1$ *to* $|V|$ **do**
    $V_{neighbor}, E_{neighbor} \leftarrow Neighbors(\mathbf{v}^{(i)})$;
    $\mathbf{v}_l^{(i)} \leftarrow \sum_{\mathbf{v}^{(j)} \in V_{neighbor}} \mathbf{v}_{l-1}^{(j)}$;
    $\mathbf{e}_l^{(i)} \leftarrow \sum_{\mathbf{e}^{(j)} \in E_{neighbor}} \mathbf{e}_0^{(j)}$;
    $\mathbf{v}_l^{(i)} \leftarrow concat(\mathbf{v}_l^{(i)}, \mathbf{e}_l^{(i)})$;
    $\mathbf{v}_l^{(i)} \leftarrow ReLU(\mathbf{v}_{l-1}^{(i)}\mathbf{U}_l + (\mathbf{v}_l^{(i)}\mathbf{W}_l)$;
    $\mathbf{v}_l^{(i)} \leftarrow softmax(\mathbf{v}_l^{(i)}\mathbf{H}_l^{|V_{neighbor}|})$;
    $\mathbf{f} \leftarrow \mathbf{f} + \mathbf{v}_l^{(i)}$;
  **end**
**end**

#### Supplementary Note 3: Learning-to-rank Objective Functions

CIGER treats the gene expression profiles as the lists ranked by their z-score values and then minimizes several learning-to-rank objective functions including both pair-wise (i.e. RankNet) and list-wise (i.e. ListNet, ListMLE, and RankCosine) functions between the predicted ($\mathbf{Y}$) and the ground-truth ($\mathbf{Z}$) gene expression profiles. The details of these learning-to-rank objective functions are presented in the following paragraphs.

**ListNet**[2] is the list-wise ranking objective function that minimizes the cross-entropy loss between top-1 probability of the predicted and ground-truth gene expression profiles. In particular, the top-1 probability of gene $i$ in the gene expression profile $j$ is the probability of that gene being ranked first among all genes in that profile and is computed as follows:

$$P_{top-1}^{z^{(j)}}(x_i^{(j)}) = \frac{\exp(z_i^{(j)})}{\sum_{k=1}^{n_g}\exp(z_k^{(j)})}$$

and then ListNet minimizes the loss:

$$L_{ListNet} = -\sum_{j=1}^{n_b}\sum_{k=1}^{n_g} P_{top-1}^{z^{(j)}}(x_i^{(j)})\log(P_{top-1}^{y^{(j)}}(x_i^{(j)}))$$

**ListMLE**[3] is the list-wise ranking objective function that maximizes the likelihood of the rank given the list of gene expression values. In particular, let $\pi^{(j)}$ is the ranked list given the gene expression profile $\mathbf{z}^{(j)}$, the negative log-likelihood of the ranked lists is computed as follows:

$$L_{ListMLE} = -\sum_{j=1}^{n_b}\sum_{i=1}^{n_g} \frac{\exp(y^{(j)}_{\pi_i^{(j)}})}{\sum_{k=i}^{n_g}\exp(y^{(j)}_{\pi_k^{(j)}})}$$

**RankCosine**[4] is the list-wise ranking objective function that measures the cosine similarity between the predicted and ground-truth ranking lists. In particular, this score is computed as follows:

$$L_{RankCosine} = \frac{1}{2}\sum_{j=1}^{n_b}\left(1 - \frac{\mathbf{z}^{(j)T}\mathbf{y}^{(j)}}{\|\mathbf{z}^{(j)}\|\,\|\mathbf{y}^{(j)}\|}\right)$$

**RankNet**[5] is the pair-wise ranking objective function that considers the ranking between every pairs of genes. In particular, given the ranked list $\pi^{(j)}$ of gene expression profiles $\mathbf{z}^{(j)}$, RankNet is computed as the cross-entropy loss between predicted and ground-truth pair-wise ranked probabilities as follows:

$$L_{RankNet} = \sum_{j=1}^{n_b}\sum_{i,k=1}^{n_g} -\hat{P}^{(j)}_{ik}\log(P^{(j)}_{ik}) - (1-\hat{P}^{(j)}_{ik})\log(1-P^{(j)}_{ik})$$

where $\hat{P}^{(j)}_{ik}$ and $P^{(j)}_{ik}$ are the ground-truth and predicted probabilities that gene $i$ is ranked higher than gene $k$ and are computed as follows:

$$\hat{P}^{(j)}_{ik} = \frac{1}{2}(1+S^{(j)}_{ik})$$

$$P^{(j)}_{ik} = \frac{1}{1+exp(-(y^{(j)}_i - y^{(j)}_k))}$$

where $S^{(j)}_{ik}$ = 1 if gene $i$ is ranked higher than gene $k$ in the ranked list $\pi^{(j)}$, -1 if gene $k$ is ranked higher, and 0.5 if they are ranked equally.

### Supplementary Tables

| Metrics | Models | A375 | A549 | HA1E | HCC515 | HELA | HT29 | MCF7 | PC3 | VCAP | YAPC |
|---|---|---|---|---|---|---|---|---|---|---|---|
| NDCG | TT-WOPT | 0.7421 ± 0.0041 | 0.7367 ± 0.0103 | 0.7423 ± 0.0035 | 0.7346 ± 0.0059 | 0.7303 ± 0.0111 | 0.7468 ± 0.0091 | 0.7415 ± 0.0036 | 0.7298 ± 0.0028 | 0.7371 ± 0.0066 | 0.7386 ± 0.0044 |
| | DeepCOP | 0.8134 ± 0.0053 | 0.7980 ± 0.0091 | 0.7957 ± 0.0087 | 0.7960 ± 0.0031 | 0.8013 ± 0.0050 | 0.8253 ± 0.0073 | 0.8140 ± 0.0016 | 0.8123 ± 0.0044 | 0.8024 ± 0.0106 | 0.8152 ± 0.0086 |
| | CIGER | **0.8357 ± 0.0067** | **0.8188 ± 0.0129** | **0.8254 ± 0.0047** | **0.8089 ± 0.0058** | **0.8332 ± 0.0169** | **0.8362 ± 0.0115** | **0.8276 ± 0.0101** | **0.8267 ± 0.0034** | **0.8298 ± 0.0073** | **0.8315 ± 0.0074** |
| P@10 | TT-WOPT | 0.2601 ± 0.0225 | 0.2476 ± 0.0179 | 0.2713 ± 0.0243 | 0.2482 ± 0.0129 | 0.2482 ± 0.0426 | 0.2790 ± 0.0419 | 0.2612 ± 0.0212 | 0.2499 ± 0.0258 | 0.2624 ± 0.0304 | 0.2813 ± 0.0556 |
| | DeepCOP | 0.5417 ± 0.0295 | 0.4935 ± 0.0502 | 0.5166 ± 0.0186 | 0.4975 ± 0.0187 | 0.5368 ± 0.0528 | 0.5865 ± 0.0201 | 0.5532 ± 0.0047 | 0.5757 ± 0.0323 | 0.5160 ± 0.0513 | 0.5913 ± 0.0447 |
| | CIGER | **0.6021 ± 0.0363** | **0.5468 ± 0.0631** | **0.5840 ± 0.0139** | **0.5365 ± 0.0292** | **0.6432 ± 0.0530** | **0.6055 ± 0.0481** | **0.5898 ± 0.0330** | **0.6232 ± 0.0164** | **0.6248 ± 0.0231** | **0.6290 ± 0.0159** |
| P@50 | TT-WOPT | 0.2423 ± 0.0178 | 0.2345 ± 0.0160 | 0.2499 ± 0.0149 | 0.2328 ± 0.0096 | 0.2301 ± 0.0332 | 0.2552 ± 0.0283 | 0.2396 ± 0.0151 | 0.2290 ± 0.0211 | 0.2363 ± 0.0214 | 0.2427 ± 0.0354 |
| | DeepCOP | 0.4757 ± 0.0176 | 0.4373 ± 0.0428 | 0.4250 ± 0.0211 | 0.4267 ± 0.0136 | 0.4558 ± 0.0253 | 0.5122 ± 0.0179 | 0.4654 ± 0.0090 | 0.4951 ± 0.0246 | 0.4486 ± 0.0376 | 0.5072 ± 0.0301 |
| | CIGER | **0.5420 ± 0.0302** | **0.4994 ± 0.0453** | **0.5134 ± 0.0055** | **0.4768 ± 0.0228** | **0.5578 ± 0.0509** | **0.5421 ± 0.0328** | **0.5191 ± 0.0355** | **0.5451 ± 0.0188** | **0.5435 ± 0.0188** | **0.5408 ± 0.0141** |
| P@100 | TT-WOPT | 0.2316 ± 0.0138 | 0.2249 ± 0.0194 | 0.2368 ± 0.0142 | 0.2229 ± 0.0039 | 0.2185 ± 0.0252 | 0.2369 ± 0.0235 | 0.2289 ± 0.0121 | 0.2193 ± 0.0137 | 0.2295 ± 0.0164 | 0.2321 ± 0.0290 |
| | DeepCOP | 0.4262 ± 0.0140 | 0.3902 ± 0.0345 | 0.3705 ± 0.0182 | 0.3856 ± 0.0072 | 0.4051 ± 0.0187 | 0.4613 ± 0.0185 | 0.4197 ± 0.0107 | 0.4418 ± 0.0193 | 0.4083 ± 0.0259 | 0.4395 ± 0.0215 |
| | CIGER | **0.4877 ± 0.0239** | **0.4545 ± 0.0380** | **0.4632 ± 0.0052** | **0.4324 ± 0.0160** | **0.4984 ± 0.0490** | **0.4898 ± 0.0288** | **0.4672 ± 0.0321** | **0.4851 ± 0.0125** | **0.4757 ± 0.0131** | **0.4806 ± 0.0131** |
| P@200 | TT-WOPT | 0.2211 ± 0.0114 | 0.2167 ± 0.0154 | 0.2258 ± 0.0104 | 0.2122 ± 0.0048 | 0.2087 ± 0.0184 | 0.2224 ± 0.0172 | 0.2193 ± 0.0099 | 0.2108 ± 0.0109 | 0.2185 ± 0.0127 | 0.2219 ± 0.0225 |
| | DeepCOP | 0.3646 ± 0.0093 | 0.3389 ± 0.0228 | 0.3207 ± 0.0161 | 0.3339 ± 0.0068 | 0.3462 ± 0.0147 | 0.3931 ± 0.0129 | 0.3603 ± 0.0055 | 0.3722 ± 0.0119 | 0.3460 ± 0.0143 | 0.3687 ± 0.0150 |
| | CIGER | **0.4172 ± 0.0180** | **0.3879 ± 0.0252** | **0.3991 ± 0.0031** | **0.3716 ± 0.0134** | **0.4163 ± 0.0385** | **0.4174 ± 0.0219** | **0.3973 ± 0.0240** | **0.4070 ± 0.0085** | **0.3998 ± 0.0081** | **0.4119 ± 0.0072** |

**Supplementary Table S1.** Cell-specific performances (NDCG and Precision@K) of TT-WOPT, DeepCOP, and CIGER for up-regulated gene ranking under 5-fold cross-validation setting.

| Metrics | Models | A375 | A549 | HA1E | HCC515 | HELA | HT29 | MCF7 | PC3 | VCAP | YAPC |
|---|---|---|---|---|---|---|---|---|---|---|---|
| NDCG | TT-WOPT | 0.7625 ± 0.0033 | 0.7500 ± 0.0069 | 0.7583 ± 0.0043 | 0.7513 ± 0.0065 | 0.7500 ± 0.0084 | 0.7584 ± 0.0096 | 0.7477 ± 0.0038 | 0.7468 ± 0.0030 | 0.7508 ± 0.0083 | 0.7629 ± 0.0140 |
| | DeepCOP | 0.8441 ± 0.0032 | 0.8212 ± 0.0092 | 0.8258 ± 0.0051 | 0.8202 ± 0.0123 | 0.8173 ± 0.0039 | 0.8372 ± 0.0047 | 0.8245 ± 0.0058 | 0.8278 ± 0.0053 | 0.8394 ± 0.0073 | 0.8470 ± 0.0037 |
| | CIGER | **0.8574 ± 0.0046** | **0.8378 ± 0.0096** | **0.8495 ± 0.0044** | **0.8251 ± 0.0087** | **0.8552 ± 0.0136** | **0.8519 ± 0.0053** | **0.8346 ± 0.0091** | **0.8412 ± 0.0015** | **0.8599 ± 0.0089** | **0.8625 ± 0.0028** |
| P@10 | TT-WOPT | 0.2916 ± 0.0093 | 0.2941 ± 0.0172 | 0.2984 ± 0.0072 | 0.2782 ± 0.0414 | 0.2657 ± 0.0499 | 0.3214 ± 0.0386 | 0.2806 ± 0.0160 | 0.2683 ± 0.0169 | 0.2893 ± 0.0298 | 0.2909 ± 0.0372 |
| | DeepCOP | 0.6355 ± 0.0167 | 0.5473 ± 0.0625 | 0.5334 ± 0.0288 | 0.5262 ± 0.0566 | 0.5316 ± 0.0612 | 0.6052 ± 0.0333 | 0.5883 ± 0.0198 | 0.5793 ± 0.0353 | 0.6178 ± 0.0379 | 0.6368 ± 0.0202 |
| | CIGER | **0.6578 ± 0.0223** | **0.6151 ± 0.0611** | **0.6365 ± 0.0181** | **0.5472 ± 0.0329** | **0.6885 ± 0.0414** | **0.6522 ± 0.0258** | **0.6069 ± 0.0349** | **0.6198 ± 0.0201** | **0.6788 ± 0.0360** | **0.7004 ± 0.0520** |
| P@50 | TT-WOPT | 0.2708 ± 0.0032 | 0.2615 ± 0.0169 | 0.2653 ± 0.0117 | 0.2553 ± 0.0231 | 0.2480 ± 0.0362 | 0.2800 ± 0.0285 | 0.2611 ± 0.0111 | 0.2462 ± 0.0157 | 0.2674 ± 0.0305 | 0.2647 ± 0.0278 |
| | DeepCOP | 0.5473 ± 0.0163 | 0.4946 ± 0.0415 | 0.4748 ± 0.0230 | 0.4740 ± 0.0409 | 0.4652 ± 0.0234 | 0.5489 ± 0.0291 | 0.5157 ± 0.0160 | 0.5177 ± 0.0269 | 0.5479 ± 0.0317 | 0.5561 ± 0.0153 |
| | CIGER | **0.5983 ± 0.0104** | **0.5523 ± 0.0436** | **0.5715 ± 0.0131** | **0.5038 ± 0.0267** | **0.6025 ± 0.0440** | **0.6019 ± 0.0220** | **0.5524 ± 0.0274** | **0.5655 ± 0.0074** | **0.6269 ± 0.0346** | **0.6162 ± 0.0132** |
| P@100 | TT-WOPT | 0.2562 ± 0.0034 | 0.2452 ± 0.0189 | 0.2494 ± 0.0092 | 0.2398 ± 0.0157 | 0.2309 ± 0.0285 | 0.2614 ± 0.0234 | 0.2481 ± 0.0133 | 0.2348 ± 0.0113 | 0.2503 ± 0.0251 | 0.2528 ± 0.0269 |
| | DeepCOP | 0.4890 ± 0.0127 | 0.4454 ± 0.0350 | 0.4348 ± 0.0219 | 0.4347 ± 0.0355 | 0.4213 ± 0.0164 | 0.4950 ± 0.0233 | 0.4648 ± 0.0125 | 0.4713 ± 0.0196 | 0.4925 ± 0.0269 | 0.5001 ± 0.0146 |
| | CIGER | **0.5425 ± 0.0100** | **0.5078 ± 0.0344** | **0.5218 ± 0.0126** | **0.4652 ± 0.0243** | **0.5392 ± 0.0389** | **0.5530 ± 0.0226** | **0.5026 ± 0.0215** | **0.5183 ± 0.0045** | **0.5770 ± 0.0311** | **0.5543 ± 0.0114** |
| P@200 | TT-WOPT | 0.2358 ± 0.0068 | 0.2300 ± 0.0154 | 0.2324 ± 0.0084 | 0.2239 ± 0.0068 | 0.2189 ± 0.0225 | 0.2387 ± 0.0165 | 0.2304 ± 0.0110 | 0.2205 ± 0.0075 | 0.2318 ± 0.0188 | 0.2337 ± 0.0178 |
| | DeepCOP | 0.4144 ± 0.0080 | 0.3839 ± 0.0253 | 0.3855 ± 0.0155 | 0.3771 ± 0.0260 | 0.3706 ± 0.0101 | 0.4194 ± 0.0164 | 0.3985 ± 0.0101 | 0.4049 ± 0.0138 | 0.4214 ± 0.0170 | 0.4280 ± 0.0140 |
| | CIGER | **0.4609 ± 0.0080** | **0.4273 ± 0.0267** | **0.4465 ± 0.0098** | **0.3985 ± 0.0169** | **0.4589 ± 0.0320** | **0.4680 ± 0.0161** | **0.4286 ± 0.0170** | **0.4422 ± 0.0046** | **0.4894 ± 0.0223** | **0.4657 ± 0.0084** |

**Supplementary Table S2.** Cell-specific performances (NDCG and Precision@K) of TT-WOPT, DeepCOP, and CIGER for down-regulated gene ranking under 5-fold cross-validation setting.

| Model | Classification Task | | | |
|---|---|---|---|---|
| | Up-regulated | | Down-regulated | |
| | AU-PRC | F1 | AU-PRC | F1 |
| TT-WOPT | 0.0527 ± 0.0010 | 0.0953 ± 0.0038 | 0.0563 ± 0.0013 | 0.0977 ± 0.0025 |
| Logistic Regression | 0.0848 ± 0.0065 | 0.1437 ± 0.0093 | 0.0961 ± 0.0053 | 0.1588 ± 0.0070 |
| DeepCOP | 0.1137 ± 0.0096 | 0.1733 ± 0.0078 | 0.1250 ± 0.0114 | 0.1894 ± 0.0126 |
| CIGER | **0.1498** ± **0.0052** | **0.2226** ± **0.0076** | **0.1542** ± **0.0082** | **0.2335** ± **0.0083** |

**Supplementary Table S3.** Performances (i.e., AU-PRC and F1) of TT-WOPT, Logistic Regression, DeepCOP, and CIGER for up-regulated and down-regulated gene classification under 5-fold cross-validation setting.

| Up-regulated gene ranking | | | | | |
|---|---|---|---|---|---|
| Setting | NDCG | P@10 | P@50 | P@100 | P@200 |
| Full data | 0.7761 ± 0.0029 | 0.4070 ± 0.0028 | 0.3447 ± 0.0042 | 0.3124 ± 0.0085 | 0.2804 ± 0.0091 |
| Filtered data | **0.8275** ± **0.0041** | **0.5973** ± **0.0170** | **0.5276** ± **0.0126** | **0.4735** ± **0.0101** | **0.4027** ± **0.0077** |
| Down-regulated gene ranking | | | | | |
| Setting | NDCG | P@10 | P@50 | P@100 | P@200 |
| Full data | 0.7966 ± 0.0049 | 0.4762 ± 0.0164 | 0.4079 ± 0.0173 | 0.3281 ± 0.0235 | 0.3182 ± 0.0136 |
| Filtered data | **0.8460** ± **0.0023** | **0.6342** ± **0.0120** | **0.5753** ± **0.0041** | **0.5250** ± **0.0034** | **0.4465** ± **0.0035** |

**Supplementary Table S4.** Average performances (NDCG and Precision@K) of CIGER when training with all gene expression profiles (i.e., full data) and high-quality gene expression profiles (i.e., filtered data) for ranking up-regulated and down-regulated genes under the 5-fold cross-validation setting.

| | A375 | | A549 | | HA1E | | HCC515 | | HELA | | HT29 | | MCF7 | | PC3 | | VCAP | | YAPC | |
|---|---|---|---|---|---|---|---|---|---|---|---|---|---|---|---|---|---|---|---|---|
| | Rank | P@200 | Rank | P@200 | Rank | P@200 | Rank | P@200 | Rank | P@200 | Rank | P@200 | Rank | P@200 | Rank | P@200 | Rank | P@200 | Rank | P@200 |
| Sucralfate | **4** | **0.3150** | **3** | **0.3300** | 14 | 0.3200 | **4** | **0.3400** | **3** | **0.3350** | **5** | **0.3700** | 11 | 0.3300 | **3** | **0.3250** | **8** | **0.3300** | **4** | **0.3400** |
| Inositol Hexasul-phate | **6** | **0.3000** | **4** | **0.3200** | 13 | 0.3200 | **3** | **0.3500** | 17 | 0.3250 | **1** | **0.3900** | **3** | **0.3650** | **4** | **0.3200** | 19 | 0.3100 | **3** | **0.3450** |
| Ginsenoside B2 | **7** | **0.3000** | **7** | **0.3100** | **2** | **0.3500** | **6** | **0.3300** | **4** | **0.3350** | **4** | **0.3750** | **7** | **0.3400** | **5** | **0.3150** | **9** | **0.3300** | **5** | **0.3350** |
| Madecassoside | **9** | **0.2850** | 11 | 0.3000 | **6** | **0.3300** | 12 | 0.3250 | **5** | **0.3350** | 13 | 0.3500 | 14 | 0.3250 | 20 | 0.2800 | 15 | 0.3200 | **6** | **0.3200** |
| Ginsenoside Rb1 | 13 | 0.2800 | **10** | **0.3000** | **3** | **0.3400** | **9** | **0.3250** | 12 | 0.3300 | 11 | 0.3550 | 15 | 0.3250 | 13 | 0.2900 | 11 | 0.3250 | 20 | 0.3100 |
| Chromium gluconate | 24 | 0.2400 | 15 | 0.2900 | **5** | **0.3350** | **10** | **0.3250** | 11 | 0.3300 | 16 | 0.3500 | 12 | 0.3250 | 15 | 0.2850 | **10** | **0.3250** | 15 | 0.3100 |
| Sodium ferric glu-conate complex | **5** | **0.3150** | **5** | **0.3100** | **9** | **0.3300** | 13 | 0.3250 | **9** | **0.3300** | 28 | 0.3350 | **9** | **0.3300** | **6** | **0.3050** | 12 | 0.3200 | 26 | 0.3000 |
| Betadex | **10** | **0.2800** | 19 | 0.2850 | 19 | 0.3050 | 32 | 0.3000 | **10** | **0.3300** | 39 | 0.3250 | 13 | 0.3250 | **8** | **0.3050** | **5** | **0.3350** | **10** | **0.3150** |
| Sucrosofate | **3** | **0.3200** | **2** | **0.3450** | 11 | 0.3250 | **1** | **0.3700** | **2** | **0.3500** | **2** | **0.3900** | **1** | **0.3700** | **2** | **0.3400** | **3** | **0.3450** | **2** | **0.3500** |

**Supplementary Table S5.** Cell-specific ranks and the corresponding Precision@200 scores of pancreatic cancer's drug candidates calculated from our drug repurposing pipeline. Drugs in top 10 cell-specific evaluations are highlighted.

| | A375 | | A549 | | HA1E | | HCC515 | | HELA | | HT29 | | MCF7 | | PC3 | | VCAP | | YAPC | |
|---|---|---|---|---|---|---|---|---|---|---|---|---|---|---|---|---|---|---|---|---|
| | Rank | GSEA | Rank | GSEA | Rank | GSEA | Rank | GSEA | Rank | GSEA | Rank | GSEA | Rank | GSEA | Rank | GSEA | Rank | GSEA | Rank | GSEA |
| Dipyridamole | **2** | **0.3475** | 3198 | 0.3021 | **1** | **0.3565** | 72 | 0.3388 | 119 | 0.2667 | 134 | 0.3270 | 23 | 0.2868 | **1** | **0.3921** | **5** | **0.4107** | **1** | **0.3657** |
| Gedatolisib | 886 | 0.0000 | 4464 | 0.0000 | 496 | 0.2597 | 59 | 0.3429 | 183 | 0.2480 | **2** | **0.3626** | 14 | 0.3006 | **2** | **0.3640** | **8** | **0.4027** | **5** | **0.3449** |
| AZD-8055 | 886 | 0.0000 | 3732 | 0.2817 | 524 | 0.2565 | **8** | **0.3770** | 98 | 0.2716 | 1891 | 0.2684 | **8** | **0.3108** | 35 | 0.3098 | 11 | 0.4006 | 72 | 0.2935 |
| Linagliptin | 886 | 0.0000 | 10374 | 0.0000 | 620 | 0.2452 | 21 | 0.3626 | 206 | 0.2415 | 1050 | 0.2916 | 21 | 0.2946 | **7** | **0.3272** | 36 | 0.3756 | **6** | **0.3386** |
| ZSTK-474 | 814 | 0.1893 | 4168 | 0.2498 | 435 | 0.2704 | **7** | **0.3773** | 181 | 0.2483 | 1006 | 0.2936 | 34 | 0.2774 | 37 | 0.3091 | **2** | **0.4214** | **3** | **0.3524** |
| Biguanide | 886 | 0.0000 | **2** | **0.4528** | 222 | 0.2882 | 2771 | 0.0000 | 687 | 0.1912 | **3** | **0.3604** | 946 | 0.0000 | 606 | 0.0000 | 200 | 0.3182 | 791 | 0.0000 |
| CH-5132799 | 886 | 0.0000 | 3503 | 0.2914 | 880 | 0.2294 | **6** | **0.3775** | 81 | 0.2771 | 1511 | 0.2776 | **10** | **0.3054** | 117 | 0.2863 | 49 | 0.3715 | 73 | 0.2929 |
| Vistusertib | 886 | 0.0000 | 3627 | 0.2868 | 532 | 0.2555 | **5** | **0.3778** | 107 | 0.2701 | 1508 | 0.2776 | **6** | **0.3185** | 49 | 0.3049 | **9** | **0.4012** | 90 | 0.2889 |
| Preladenant | 886 | 0.0000 | 3988 | 0.2658 | 504 | 0.2589 | 94 | 0.3308 | 126 | 0.2650 | 656 | 0.3091 | **7** | **0.3154** | **5** | **0.3342** | 16 | 0.3935 | **9** | **0.3338** |

**Supplementary Table S6.** Cell-specific ranks and the corresponding GSEA scores of pancreatic cancer's drug candidates calculated from our drug repurposing pipeline. Drugs in top 10 cell-specific evaluations are highlighted.

### Supplementary Figures

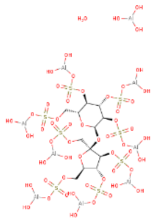

DB00364

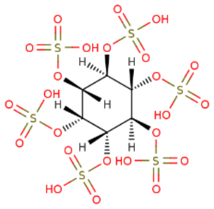

DB01666

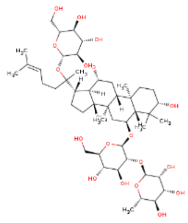

DB14815

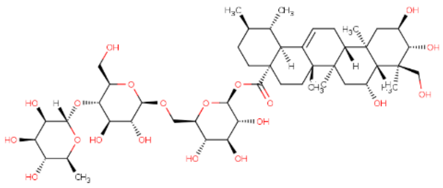

DB15532

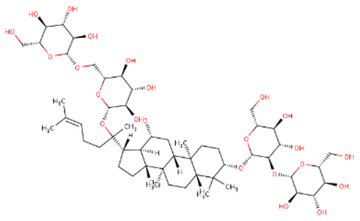

DB06749

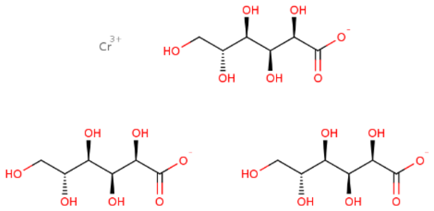

DB14528

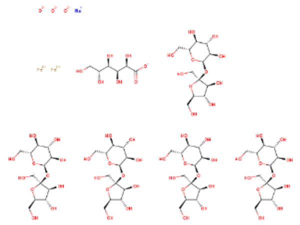

DB09517

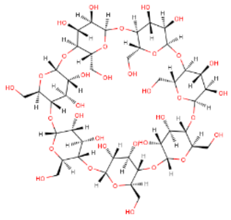

DB03995

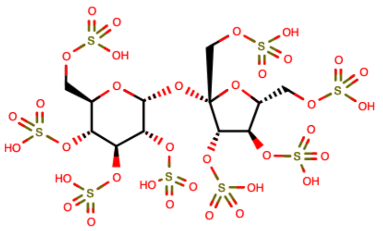

DB01901

**Supplementary Figure S1.** Drug screening pipeline using CIGER. This model is trained with LINCS L1000 dataset to learn the relation between gene expression profiles and molecular structures (i.e., SMILES). Then molecular structures retrieved from DrugBank database are putted into CIGER to generate the corresponding gene expression profiles. Finally, these profiles are compared with disease profiles calculated from treated and untreated samples to find the most potential treatments for that disease.

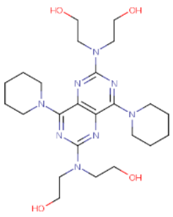

DB00975

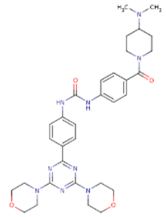

DB11896

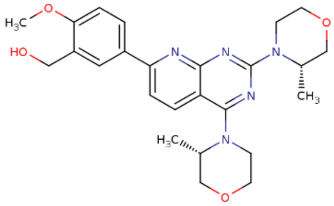

DB12774

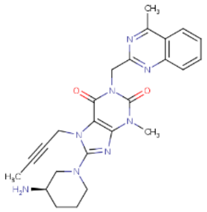

DB08882

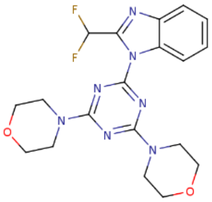

DB12904

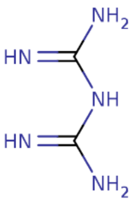

DB13100

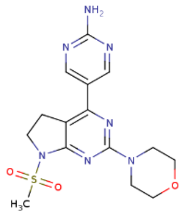

DB13051

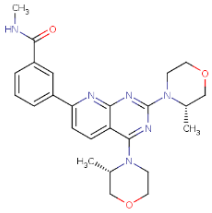

DB11925

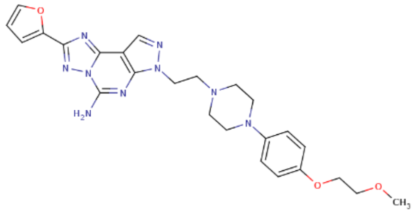

DB11864

**Supplementary Figure S2.** Drug screening pipeline using CIGER. This model is trained with LINCS L1000 dataset to learn the relation between gene expression profiles and molecular structures (i.e., SMILES). Then molecular structures retrieved from DrugBank database are putted into CIGER to generate the corresponding gene expression profiles. Finally, these profiles are compared with disease profiles calculated from treated and untreated samples to find the most potential treatments for that disease.

### References


**1.** Duvenaud, D. K. *et al.* Convolutional networks on graphs for learning molecular fingerprints. In *Advances in Neural Information Processing Systems*, 2224–2232 (2015).

**2.** Cao, Z., Qin, T., Liu, T.-Y., Tsai, M.-F. & Li, H. Learning to rank: from pairwise approach to listwise approach. In *Proceedings of the 24th International Conference on Machine Learning*, 129–136 (2007).

**3.** Xia, F., Liu, T.-Y., Wang, J., Zhang, W. & Li, H. Listwise approach to learning to rank: theory and algorithm. In *Proceedings of the 25th International Conference on Machine Learning*, 1192–1199 (2008).

**4.** Qin, T. *et al.* Query-level loss functions for information retrieval. *Inf. Process. & Manag.* **44**, 838–855 (2008).

**5.** Burges, C. *et al.* Learning to rank using gradient descent. In *Proceedings of the 22nd International Conference on Machine Learning*, 89–96 (2005).
